# The cytoplasmic tail of myelin protein zero induces morphological changes in lipid membranes

**DOI:** 10.1101/2024.03.22.586069

**Authors:** Oda C. Krokengen, Christine Touma, Anna Mularski, Aleksi Sutinen, Ryan Dunkel, Marie Ytterdal, Arne Raasakka, Haydyn D.T. Mertens, Adam Cohen Simonsen, Petri Kursula

**Affiliations:** Department of Biomedicine, University of Bergen, Bergen, Norway; Faculty of Biochemistry and Molecular Medicine & Biocenter Oulu, University of Oulu, Oulu, Finland; Department of Physics, Chemistry and Pharmacy, University of Southern Denmark, Odense, Denmark; European Molecular Biology Laboratory EMBL, Hamburg Site, c/o DESY, Hamburg, Germany

## Abstract

The major myelin protein expressed by the peripheral nervous system Schwann cells is protein zero (P0), representing 50% of the total protein content in myelin. This 30-kDa integral membrane protein consists of an immunoglobulin (Ig)-like domain, a transmembrane helix, and a 69-residue C-terminal cytoplasmic tail (P0ct). The basic residues in P0ct contribute to the tight packing of the myelin lipid bilayers, and alterations in the tail affect how P0 functions as an adhesion molecule necessary for the stability of compact myelin. Several neurodegenerative neuropathies are related to P0, including the more common Charcot-Marie-Tooth disease (CMT) and Dejerine-Sottas syndrome (DSS), but also rare cases of motor and sensory polyneuropathy. We find that high P0ct concentrations affect the membrane properties of bicelles and induce a lamellar-to-inverted hexagonal phase transition, which causes the bicelles to fuse into long, protein-containing filament-like structures. These structures likely reflect the formation of semi-crystalline lipid domains of potential relevance for myelination. Not only is P0ct important for stacking lipid membranes, but time-lapse fluorescence microscopy shows that it might affect membrane properties during myelination. We further describe recombinant production and low-resolution structural characterization of full-length human P0. Our findings shed light on P0ct effects on membrane properties, and with successful purification of full-length P0, we have new tools to study *in vitro* the role P0 has in myelin formation and maintenance.

## Introduction

The myelin sheath is the electrical insulator of our nerves. This firmly wrapped multi-layered proteolipid membrane can generate conduction velocities up to 150 m/s in myelinated axons (Jean Harry & Toews, 1998; Purves et al., 2001). Integral and peripheral membrane proteins found in myelin that can stack lipid bilayers include proteolipid protein (PLP) (Ruskamo et al., 2022), peripheral myelin protein 22 (PMP22) (Mittendorf et al., 2017), myelin basic protein (MBP) (Min et al., 2009; Raasakka et al., 2017; Suresh et al., 2010), myelin protein zero (P0) (Filbin et al., 1999; Raasakka et al., 2019b; Raasakka et al., 2019c; Wong & Filbin, 1994, 1996), and peripheral myelin protein 2 (P2) (Ruskamo et al., 2020; Sedzik et al., 1985; Uusitalo et al., 2021). P0 corresponds to 50% of total myelin protein in the peripheral nervous system (PNS), and various point mutations in human P0 result in demyelinating peripheral neuropathies, such as Charcot-Marie-Tooth disease (CMT) type 1B, Dejerine-Sottas syndrome, and congenital hypomyelination (D’Urso et al., 1999; Laulumaa S, 2018).

P0 is the only PNS myelin glycoprotein to span and compact both the major dense line (MDL; cytoplasmic apposition) and the intraperiod line (IPL; extracellular apposition). The *shiverer* mouse, which carries a spontaneous MBP knockout, maintains a functionally normal PNS (Inouye et al., 1985). This suggests that P0 can maintain myelin compaction in the absence of MBP. P0 is not only required during the outset of myelin formation but in myelin maintenance at a later age, and it is sufficient for myelin compaction (Martini et al., 1995). P0 has a low expression level in Schwann cells that do not myelinate compared to myelinating Schwann cells (Jessen & Mirsky, 1999), and in non-neuronal cells overexpressing P0, cell-cell adhesion can be observed (D’Urso et al., 1990). Thus, P0 is integral in myelin formation and, consequently, the optimal functioning of the nervous system.

Since the discovery of P0, there has been little understanding of the details of its molecular involvement in myelin compaction. This 30-kDa transmembrane (TM) protein consists of an immunoglobulin (Ig)-like domain, a single TM helix, and a 69-residue C-terminal cytoplasmic tail (P0ct) (Liu et al., 2012). The extracellular domain is suggested to homotypically interact in a ‘head-to-head’ arrangement, and an N-linked oligosaccharide at position 122 is hypothesised to be responsible for the correct orientation of the entire TM glycoprotein, such that the P0 extracellular domains are optimally positioned to dimerise within the IPL (Quarles, 2002). Cryo-EM experiments on reconstituted full-length P0 isolated from bovine nerves revealed a zipper-like arrangement of the Ig-like domains at the extracellular apposition (Raasakka et al., 2019c), providing a low-resolution model for the PNS IPL. The basic residues in P0ct contribute to the tight packing of the inner leaflets of myelin lipid bilayers, and alterations in P0ct affect how P0 functions as an adhesion molecule in compact myelin (Wong & Filbin, 1994, 1996).

Human P0ct carries 21 basic and 6 acidic residues, giving it a strong positive charge (+15) (Lemke & Axel, 1985). This short segment mediates the compaction of the Schwann cell cytoplasm and contributes to the formation of the PNS myelin MDL (Shy, 2006). P0 contains three cysteine residues; Cys182 is conserved in P0ct, located at the junction between the TM domain and the tail (Bizzozero et al., 1994). Cys182 undergoes palmitoylation and is important for P0 adhesiveness. A mutation at this site resulted in loss of membrane binding and protein stability (Bharadwaj & Bizzozero, 1995; Gao et al., 2000; Wong & Filbin, 1996). Most CMT mutations found in P0 affect the Ig-like domain and the protein polarity, consequently affecting myelin morphology and Schwann cell-axonal interactions (Mandich et al., 2009; Shy, 2006).

Disease mutations in P0ct have also been identified (Choi et al., 2004; Fabrizi et al., 2006; Miltenberger-Miltenyi et al., 2009; Planté-Bordeneuve et al., 2001; Schneider-Gold et al., 2010; Shy et al., 2004; Street et al., 2002; Su et al., 1993), making this highly disordered region important in membrane compaction. P0ct is unfolded in aqueous solution, but it gains secondary structure upon lipid interactions. It was first suggested to fold into β-strands, but more recent studies strongly suggest a partly α-helical conformation for membrane-bound P0ct (Luo et al., 2007; Raasakka et al., 2019b; Raasakka et al., 2019c). In solution X-ray diffraction studies, Bragg peaks emerge in a concentration-dependent manner when P0ct is mixed with small unilamellar lipid vesicles (Raasakka et al., 2019c), indicating P0ct-induced formation of ordered membrane multilayers with a well-defined repeat distance.

Different membrane-mimicking model systems exist for investigating membrane protein structure and function in a lipid environment. For many years, these model membranes have been predominantly monolayers and liposomes (Akbarzadeh et al., 2013; Chan & Boxer, 2007; Eeman & Deleu, 2010; Sharma & Sharma, 1997), but bicelles have been increasingly used for studying membrane proteins. Bicelles were originally developed for NMR studies (Prosser et al., 2006), but have been adapted to other methods, as they may bridge the gap between liposomes and the native flat bilayers found in organisms. While liposomes are spherical, bicelles are flat discs with a rim of detergent (Dürr et al., 2013). Such nano-membranes are important tools to study the functions and properties of membrane proteins, as information can be extracted using different techniques, such as X-ray crystallography (Murugova et al., 2022; Ujwal & Bowie, 2011), cryo-EM (Wang & Sigworth, 2010; Yao et al., 2020), small-angle X-ray and neutron scattering (Conn et al., 2020; Dos Santos Morais et al., 2017), atomic force microscopy (Borrell et al., 2015; Sebinelli et al., 2019), and X-ray diffraction (Ruskamo et al., 2020).

Here, we further investigated the function of P0ct, while also examining the implications of using bicelles as a myelin-mimetic membrane. Expanding on our past studies using liposomes, the use of bicelles impacts the protein-lipid interactions, giving us new information about the dynamics of P0ct in the myelin sheath. We report the first instance of full-length human P0 (hP0) being expressed and purified recombinantly from insect cells. We present a low-resolution structural model for full-length hP0 in detergent micelles. The data provide novel insights into myelin protein-lipid interactions and give starting points for new lines of study, including myelin protein-induced lipid phase transitions and high-resolution structure of full-length P0 and its disease variants.

## Materials and methods

### Expression and purification of P0ct

P0ct was expressed as a fusion with an N-terminal His_6_ tag and maltose-binding protein (MaBP). Cloning, expression, and purification of His-MaBP-P0ct have been described earlier (Raasakka et al., 2019c). Briefly, His-MaBP-P0ct was expressed in *Escherichia coli* BL21(DE3) using 0.4 mM IPTG induction for 3 h in LB medium containing 100 μg/ml ampicillin, at 37 °C and 180 rpm shaking. After expression, the cells were collected by centrifugation and resuspended in 40 mM Tris-HCl, 400 mM NaCl, 20 mM imidazole, pH 8.5 with the addition of EDTA-free protease inhibitor cocktail (Roche). Cells were then lysed using ultrasonication and centrifuged to clarity, and the supernatant was collected before continuing the purification by Ni-NTA and amylose affinity chromatography. Elution from the Ni-NTA affinity matrix was performed using 32 mM HEPES (pH 7.5), 320 mM NaCl, 500 mM imidazole, before dialysis against 20 mM Tris-HCl, 400 mM NaCl, 1 mM DTT, pH 8.5, at +4 °C. Recombinant TEV protease was used for His-MaBP tag removal. Cleaved P0ct was separated from the cleaved tag and uncleaved protein by amylose affinity chromatography. Size exclusion chromatography (SEC) using a HiLoad Superdex 75 pg 16/60 column (GE Healthcare) was used to separate the cleaved protein from any contaminants, resulting in a peak corresponding to pure monodisperse P0ct. SEC buffer was 20 mM HEPES, 150 mM NaCl, pH 7.5.

### Production of hP0 bacmid

Recombinant bacmid production was carried out under manufacturer’s instructions (Bac-to-bac system, Life Technologies). Competent *E. coli* DH10Bac cells were transformed with 1 ng of pFastBac Dual hP0-GFP-6xHis (Invitrogen; a TEV cleavage site exists between hP0 and its C-terminal tags). The correct isolate was selected using LB plates containing 40 µg/ml Blue-gal, 10 µg/ml tetracycline, 50 µg/ml kanamycin, 7.5 µg/ml gentamycin, 40 µg/ml IPTG, and were incubated for 48 h at 37 °C. A white colony was selected and restreaked onto a fresh plate and left overnight at 37 °C. A further single white colony was selected for growth in liquid LB culture containing the mentioned antibiotics and was used to purify the bacmid using a standard alkaline lysis process. Diagnostic PCR confirmed the transposition of the gene of interest in the bacmid.

### Expression and purification of hP0

Sf9 insect cells cultured in Insect-XPRESS media (Lonza) were transfected with 2 µg of bacmid using Fugene 6 transfection reagent (Promega), as described by the manufacturer. The medium was supplemented with 10% FCS, 10 Units/ml penicillin, and 10 µ/ml streptomycin, and cells were cultured for 7 days at 300 rpm, 27 °C. First-generation baculovirus was harvested by centrifugation and used to infect fresh Sf9 cells in a 1 l suspension culture to induce expression over 3 days at 300 rpm, 27 °C. Expression was monitored using the fluorescence properties of GFP; its excitation peak is at 475 nm and emission peak at 511 nm. The hP0-expressing cells were washed with PBS, harvested by centrifugation, flash-frozen in liquid N_2_, and stored at −80 °C.

The cell pellets were resuspended in wash buffer (20 mM Tris pH 8.0, 500 mM NaCl, 1 mM TCEP, 50 mM imidazole) and a douncer was used to rupture the cells. The lysate was ultracentrifuged at 235 000 g for 1.5 h, the resulting pellet containing membrane material was resuspended in wash buffer, and a douncer was used to homogenise. The membrane homogenate was incubated at 4 °C with stirring overnight and was supplemented with 1% (w/v) DDM, 20 µg/ml DNase I, 10 mM MgCl_2_, 1x protease inhibitor cocktail, and 1 mM TCEP. The homogenate was ultracentrifuged at 235 000 g for 1.5 h, and the supernatant was applied onto an equilibrated Ni-NTA slurry and allowed to bind at 4 °C for 1 h. Unbound material was removed, and the matrix was washed with wash buffer supplemented with 0.03% (w/v) DDM. The fusion protein was eluted with elution buffer (20 mM Tris pH 8.0, 150 mM NaCl, 1 mM TCEP, 500 mM imidazole, 0.03% DDM) and dialysed against SEC buffer (20 mM Tris pH 8.0, 150 mM NaCl, 1 mM TCEP, 0.03% DDM) at 4 °C overnight in the presence of TEV protease. Reverse IMAC was performed to remove TEV and intact fusion protein. The cleaved hP0 was injected into an S200 Increase 10/300 GL column (GE Healthcare), and the peak corresponding to pure monomeric hP0 was collected and concentrated using Vivaspin centrifugal concentrators (Sartorius).

### Vesicle preparation

*For all experiments except membrane patch experiments,* DMPC (dimyristoylphosphatidylcholine) and DMPG (dimyristoylphosphatidylglycerol) were purchased from Larodan Fine Chemicals AB (Malmö, Sweden). Lipid stocks were prepared by dissolving dry lipids in chloroform or chloroform:methanol (75:25) to a final concentration of 10 mg/ml. The solvent was evaporated under a stream of nitrogen gas before freeze-drying overnight at −52 °C under vacuum. The dried lipid mixture was stored air-tight at −20 °C until use.

For preparation of either liposomes or bicelles, the dried lipids were hydrated with either water or HBS (20 mM HEPES, 150 mM NaCl, pH 7.5) at a final concentration of 5-10 mg/ml, inverting at room temperature overnight to ensure that no unsuspended lipids remained in the glass vial. The formed multilamellar vesicles (MLVs) were subjected to freeze-thawing using liquid N_2_ and a warm water bath with 30 s of strong vortexing. The cycle was repeated 7 times. Small unilamellar vesicles (SUVs) were prepared by sonicating fresh MLVs using a probe tip sonicator (Branson Model 450, Sonics & Materials Inc. Vibra-cell VC-130) until the lipid suspension was clear, while avoiding overheating. Bicelles were made by adding *n*-dodecylphosphocholine (DPC) to the MLVs to a q-ratio of 2.84, while pipetting up and down until the solution became clear. The SUVs and bicelles were left at room temperature for at least a couple of hours before use.

*For membrane patch experiments*, DOPC (1,2-dioleoyl-*sn*-glycero-3-phosphocholine) and DOPS (1,2-dioleoyl-*sn*-glycero-3-phospho-*L*-serine) were purchased from Avanti Polar Lipids (Alabaster, Alabama, USA). Lipid stocks containing DOPC and DOPS were prepared by dissolving dry lipids in methanol (hypergrade for LC-MS, Merck) to a final concentration of 10 mM with a DOPC:DOPS ratio of 9:1. When the lipids were completely dissolved, 0.05% DiD-C18 probe (Thermo-Invitrogen) was added. The lipid mixture was immediately used to make the lipid films for the patch experiments.

### Membrane patch experiments

Mica substrates (Plano GmbH), glued to glass coverslips using a silicone elastomer (MED-6215, Nusil Technology), were cleaved immediately before use. 50 µl of 10 mM DOPC:DOPS 9:1 (method described above) was applied onto the mica, spun on a spin coater (KW-4A, Chemat Technology) at 150 g for 40 s and placed under vacuum in a desiccator for 10-12 h to ensure solvent evaporation. The spin-coated lipid film was hydrated in 10 mM TRIS buffer (2-amino-2-(hydroxymethyl)propane-1,3-diol), 140 mM NaCl, 2 mM Ca^2+^, pH 7.4), for ≥1 h at 60 °C before doing buffer-exchange >10 times to prepare defined secondary bilayer patches. The hydrated membrane patches were then cooled down to room temperature (22 °C) and equilibrated for at least 1 h before experiments.

The response of bilayer patches to the addition of myelin proteins was monitored with time-lapse *epi*-fluorescence microscopy using a Nikon Ti2-E inverted microscope with 40 × objective (Nikon ELWD S Plan Fluor, NA = 0.6). Fluorescence excitation at 550 nm was achieved using a white light fluorescence illumination system (CoolLED, pE-300). Emission was collected at 640 nm (Nikon Cy5-A, M376571). Images were captured at 10 fps with a digital CMOS camera (ORCA-Flash 4.0 V3, Hamamatsu, 2048 × 2044 pixels) using NIS-Elements software (Nikon). At least 3 experiments per condition were performed. For each experiment, P0ct were added to the fluid cell from a known concentrated solution, such that the final bulk concentration was 10 μM.

### Synchrotron radiation circular dichroism spectroscopy

P0ct was dialyzed into Milli-Q water before SRCD measurements to exclude absorption effects from buffer components. Protein alone was measured at 0.4 mg/ml in a 100-μm pathlength closed cylindrical cell (Suprasil, Hellma Analytics). SUVs and bicelles (DMPC:DMPG at ratios 1:1, 4:1, and 9:1) were hydrated in Milli-Q and further prepared as described above. Samples containing lipids were prepared right before measurements by mixing P0ct at 0.4 mg/ml to a final molar protein-to-lipid ratio (P/L) of 1:50, 1:100, 1:200, and 1:500 with either SUVs or bicelles. Spectra were recorded from 170 to 280 nm at 30 °C. The baseline was subtracted from each dataset before converting CD units to Δε (M^−1^ cm^−1^), using P0ct concentration determined from absorbance at 280 nm. Measurements were done several times to ensure reproducibility. SRCD data were collected on the AU-CD beamline at the ASTRID2 synchrotron (ISA, Aarhus, Denmark). Thermal melting of P0ct in solution and with two different ratios of bicelles (P/L 1:50 and 1:100) was executed with temperatures ranging from 24 – 84°C at roughly 5 °C intervals. Six scans were done at each temperature with an equilibration of 5 min. After reaching the maximum temperature, a new set of 6 scans were done at 24.2 °C to check for refolding. Melting temperature (T_m_) was obtained by fitting the SRCD data at 222 nm to a sigmoidal model (GraphPad Prism 9).

### Stopped-flow SRCD

An SX-20 stopped-flow instrument (Applied Photophysics) mounted on the AU-rSRCD branch line of the AU-AMO beamline at ASTRID2 (ISA, Aarhus, Denmark) was used for data collection at 10 °C. Measurements of 0.1 mg/ml P0ct and DMPC:DMPG (1:1) bicelles at P/L of 1:100 and 1:200 were achieved using a mixer (2 ms dead time) before injecting the sample into the measurement cell (160 µl total volume, 2-mm pathlength) per shot. The CD signal (mdeg) was collected at a fixed wavelength of 195/196 nm for 5 s with 5-10 repeat shots per sample, which were averaged into a single curve. Each sample was prepared in duplicate, and water baselines were subtracted from the sample data. The data were fitted to different exponential functions using GraphPad Prism 9 (Supplementary Table 1).

### Transmission electron microscopy

P0ct was mixed with DMCP:DMPG (1:1) SUVs or bicelles in HBS to final P/L of 1:50, 1:100, and 1:200 before incubating for 1 h at room temperature. 4 µl of sample were added to a carbon-coated copper grid (300-200 mesh) before incubating for 1 min. Excess solution was removed with filter paper (Whatman), and the sample was stained with 2% uranyl acetate for 12 s. Excess liquid was removed before air-drying. TEM imaging was performed using a Jeol JEM-1230 (MedWOW) instrument. The samples were prepared in duplicate and images were obtained at several locations on the grid.

### Small-angle X-ray diffraction

SAXD experiments were performed to investigate repetitive structures. 30 μM or 60 μM P0ct were mixed with SUVs or bicelles containing 3 mM DMPC:DMPG (1:1) in HBS at room temperature and exposed to X-rays on the CoSAXS beamline, MAX IV Laboratory, Lund University (Lund, Sweden) (Kahnt et al., 2021). Lipid samples without added P0ct did not show any Bragg peaks. The energy was fixed at 12.4 keV with a wavelength of 1.0 Å. Exposure time was 200 ms, collecting 300 frames for each sample under continuous flow. Data were processed and analysed using the ATSAS package (Manalastas-Cantos et al., 2021). The peak position of momentum transfer (*s*) in P0ct-lipid samples was used to calculate repeat distances (*d*) in proteolipid structures using equation 1:

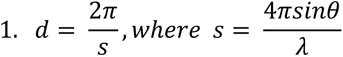

Membrane thickness of the protein-free liposomes and bicelles was determined by a modified Kratky-Porod (MKP) analysis. *Isq^4^* was plotted against *s* and fitted with a 4^th^-order polynomial, and d_m,MKP_ was calculated using equation 2 (Hägerbäumer et al., 2023):

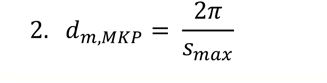

### Small-angle X-ray scattering

SEC-SAXS data were collected from 0.94 mg/ml of hP0 in 20 mM Tris pH 8, 150 mM NaCl, 1 mM TCEP, 0.03% DDM at Diamond Light Source beamline B21 (Harwell Science & Innovation Campus, Oxfordshire, United Kingdom) (Cowieson et al., 2020) and 14.4 mg/ml of P0ct in HBS on the CoSAXS beamline, MAX IV laboratory (Lund, Sweden). See Supplementary Table 2 for further details. Data collected from P0ct were processed and analysed using ATSAS (Manalastas-Cantos et al., 2021). Ensemble optimization analysis was performed using EOM (Tria et al., 2015), and *ab initio* modelling with GASBOR (Svergun et al., 2001).

The hP0 model was finalised in CORAL (Petoukhov et al., 2012). Restraints were set to the TM helix, while the N-and C-terminal domains were flexible. A hinge was present between the Ig domain and the TM, and the P0ct was built by the software. 100 DDM molecules were added around the TM helix utilizing CHARMM-GUI (Brooks et al., 2009). Visualisation was carried out in PyMOL (DeLano, 2002). AlphaFold2 (Jumper et al., 2021) was run on the Google ColabFold server (Mirdita et al., 2022) where the input was the amino acid sequence of human full-length P0 and P0ct (UniProt ID: P25189).

### DENSS electron density map modelling

Density maps were modelled in RAW 2.1.4 (Hopkins et al., 2017) using the DENSS electron density map feature (Grant, 2018). 20 reconstructions were run with the default set of 10000 electrons. Each density was aligned and averaged before a final refinement. No symmetry constraints were applied. The refined electron density map was visualised using PyMOL (DeLano, 2002).

## Results

### Liposome and bicelle characterisation

Liposomes and bicelles used to investigate P0ct were prepared from DMPC and DMPG, with DPC as detergent for the bicelles. These membrane-mimicking model systems are easily prepared, and different buffers can be applied if necessary. The membrane thickness (*d_m,MKP_*) was calculated using the MKP method (Hägerbäumer et al., 2023; Sreij et al., 2018) for samples collected at RT from several SAXS beamtimes. Following the calculations from Hägerbäumer et. al (Hägerbäumer et al., 2023), *I(q)× q^4^* was plotted against *q*, before fitting the plots with a fourth-order polynomial (Figure 1A). The SAXS curve of both model systems had the typical broad peak characteristic of a phospholipid bilayer structure. The *d_m,MKP_* was calculated from the maximal q-value using Equation (2). The sonicated liposomes have a wider span of membrane thickness than bicelles, with a mean thickness of 37.64 Å, when collecting data over several beamtimes (Figure 1B). The mean membrane thickness for the liposomes is larger than reported (Hägerbäumer et al., 2023; Sreij et al., 2018) for pure DMPC liposomes prepared by extrusion. Sonication gives are more variable size of liposomes, and the deviation is expected to be higher than in a more homogenous sample of extruded vesicles; in addition, DMPG will influence the membrane thickness. Additional variation may arise from the experimental temperature being close to that of the dimyristoyl lipid phase transition.

**Figure 1:**
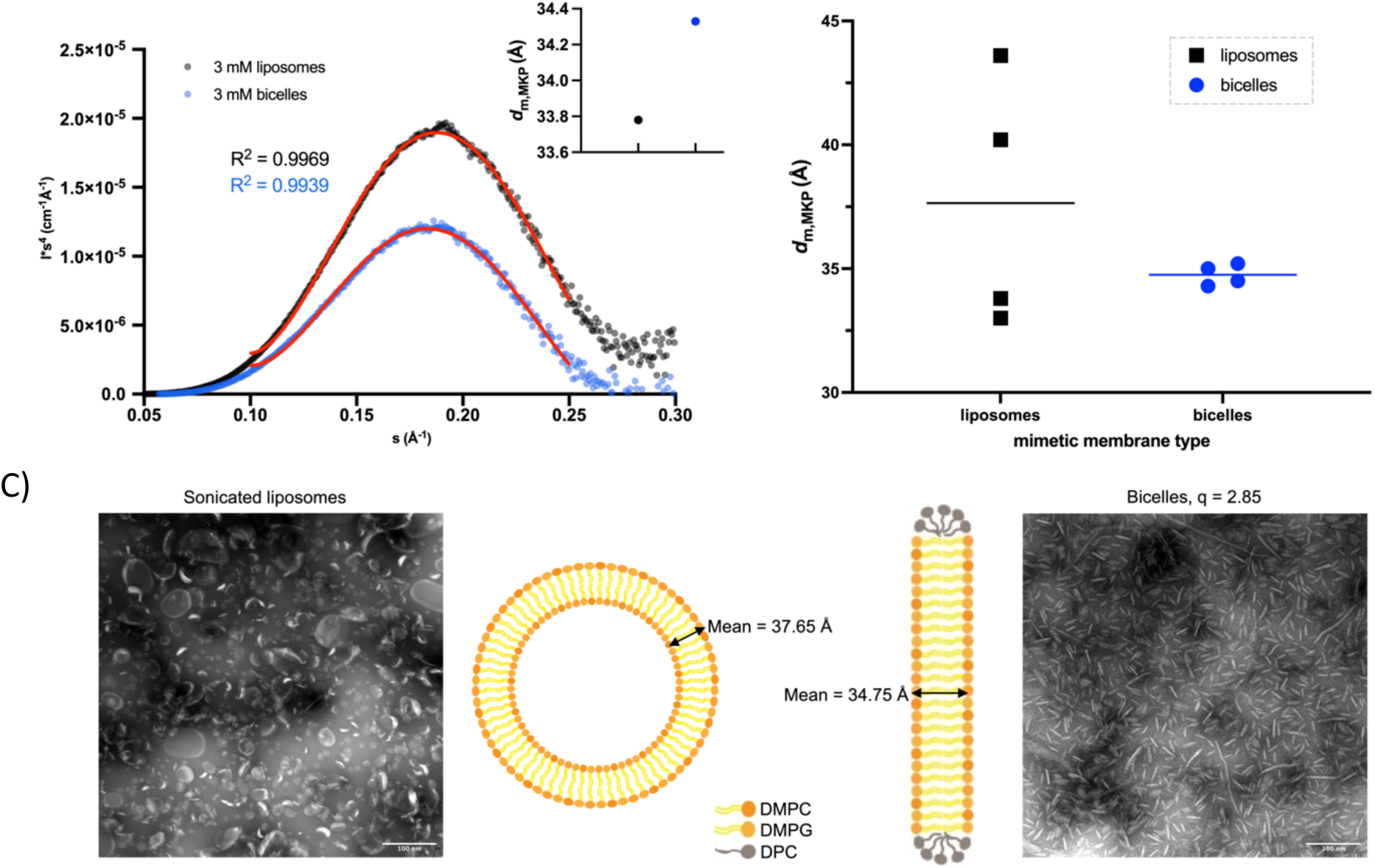
Characterisation of bicelles and sonicated liposomes. A) Membrane thickness (dm,MKP) was calculated with the MKP method (Hägerbäumer et al., 2023); I(q)×q^4^ was plotted against q before fitting the plot with a fourth-order polynomial. The maximal q-value was determined and membrane thickness calculated using equation (2), dm,MKP denoted in the upper right corner. Measurements were done at room temperature. B) dm,MKP from four separate measurements of liposomes (black) and bicelles (blue) at different time points on the MAX IV Lund CoSAXS beamline. The mean is shown as a line. C) Visualization of the two mimicking model systems using TEM with an illustration to clarify (middle). Liposomes (left) are seen as round spheres, whereas bicelles (right) are flat disks. Scale bar, 100 nm. Control TEM images of buffer and protein are shown in Supplementary Figure 1.

DMPC bicelles with 1,2-dicaproyl-*sn*-glycero-3-phosphocholine (DPCP) have a membrane thickness ranging from 36 to 40 Å with a q-ratio of 3.5 (Courreges et al., 2011), whereas bicelles with a mix of DMPC and DMPG with 3-[(3-cholamidopropyl)dimethylammonio]-1-propanesulfonate (CHAPSO) (q = 3.0) had a thickness of 51 Å (Li et al., 2013). In DMPG, the choline headgroup of DMPC is replaced by glycerol, resulting in a net negative charge. The mean membrane thickness was 34.75 Å at a q-ratio of 2.84. Q-ratio mostly determines the diameter of the bicelles (Beaugrand et al., 2014), but the thickness is also dependent on the q-ratio (Giudice et al., 2022). Moreover, doping bicelles with negatively charged lipids, DMPG in this case, will result in more rigid membranes (Katsaras et al., 2005). Consequently, using 50% DMPG might explain the decrease in membrane thickness in these bicelles compared to pure DMPC-DPCP/DHPC bicelles or DMPC:DMPG-DHPC bicelles, in which the fraction of DMPG is lower (Giudice et al., 2022). The bicelles would be highly rigid and potentially more compact.

To complement the membrane thickness analysis and to control the formation of liposomes and bicelles, we visualized the samples using TEM (shown in Figure 1C). Sonicated liposomes are expected to be spherical and to vary more in size than extruded liposomes, whereas bicelles should form when DPC is added to the MLVs-the vesicles should flatten, and the rims of the discs will be lined with detergent. The TEM images revealed polydisperse vesicles and bicelles with little to no aggregation. The results confirm the homogeneous nature of the bicelle population, in line with the above X-ray scattering analysis. Both membrane-mimicking systems resemble previously published data (Raasakka et al., 2019c; Ruskamo et al., 2020) and were further used for studying the protein-lipid interactions of P0ct.

### Secondary structure of P0ct in membrane-mimetic models

Folding investigations of P0ct have previously been done with liposomes, but not using bicelles. Earlier experiments revealed high α-helical content in various vesicle lipid mixtures and detergents (Raasakka et al., 2019b; Raasakka et al., 2019c). In the present study, the overall findings were as expected; P0ct is unfolded in water and gains helical secondary structure when interacting with the lipid models.

To investigate the effect of negatively charged lipids on the secondary structure of P0ct, various lipid mixtures were made (10%, 25%, and 50% DMPG) with P/L of 1:50, 1:100, 1:200, and 1:500 (Supplementary Figure 3). Over the various P/L ratios, the 50% DMPG liposomes and bicelles (DMPC:DMPG 1:1) seem to be most favourable for secondary structure formation with high intensity of the positive peak at 190 nm and the two negative peaks at around 208 and 220 nm. P0ct favours negatively charged lipids, and the quantity of DMPG influences the degree of folding. PNS myelin cytosolic leaflet is negatively charged, and the membrane has a high lipid-to-protein ratio (70-85% lipid by mass compared to 15-30% protein), which contributes to the close packing of the myelin sheath (Poitelon et al., 2020). This mirrors what is observed in the SRCD measurement, as high negatively charged lipid content facilitates secondary structure formation in P0ct (Supplementary Figure 3). In Figure 2A, all P/L ratios are shown with lipid mimetics containing 50% DMPG, and the difference in CD amplitude reflects the amount of aggregation of large particles, while the movement of the CD minimum peak from 200 nm towards higher wavelengths, concomitantly with a stronger band at 195 and 220 nm, is a sign of increased helical structure. The maximal level of folding seems to be reached at a P/L 1:100, and increasing the lipid amount above this level no longer induces changes in the CD peak positions. At P/L 1:50, P0ct is less folded than at P/L 1:100. Changing the model systems does not have a large effect on the secondary structure of P0ct, except for a slight increase in the intensity of the peaks when bicelles are used, most likely reflecting less aggregation. Folding of P0ct might be facilitated by flattening of the model system, potentially providing the membrane curvature needed and a larger contact area for van der Waals interactions necessary for helical folding of membrane proteins (Corin & Bowie, 2020; Corin & Bowie, 2022; Hong, 2014; Marinko et al., 2019).

**Figure 2:**
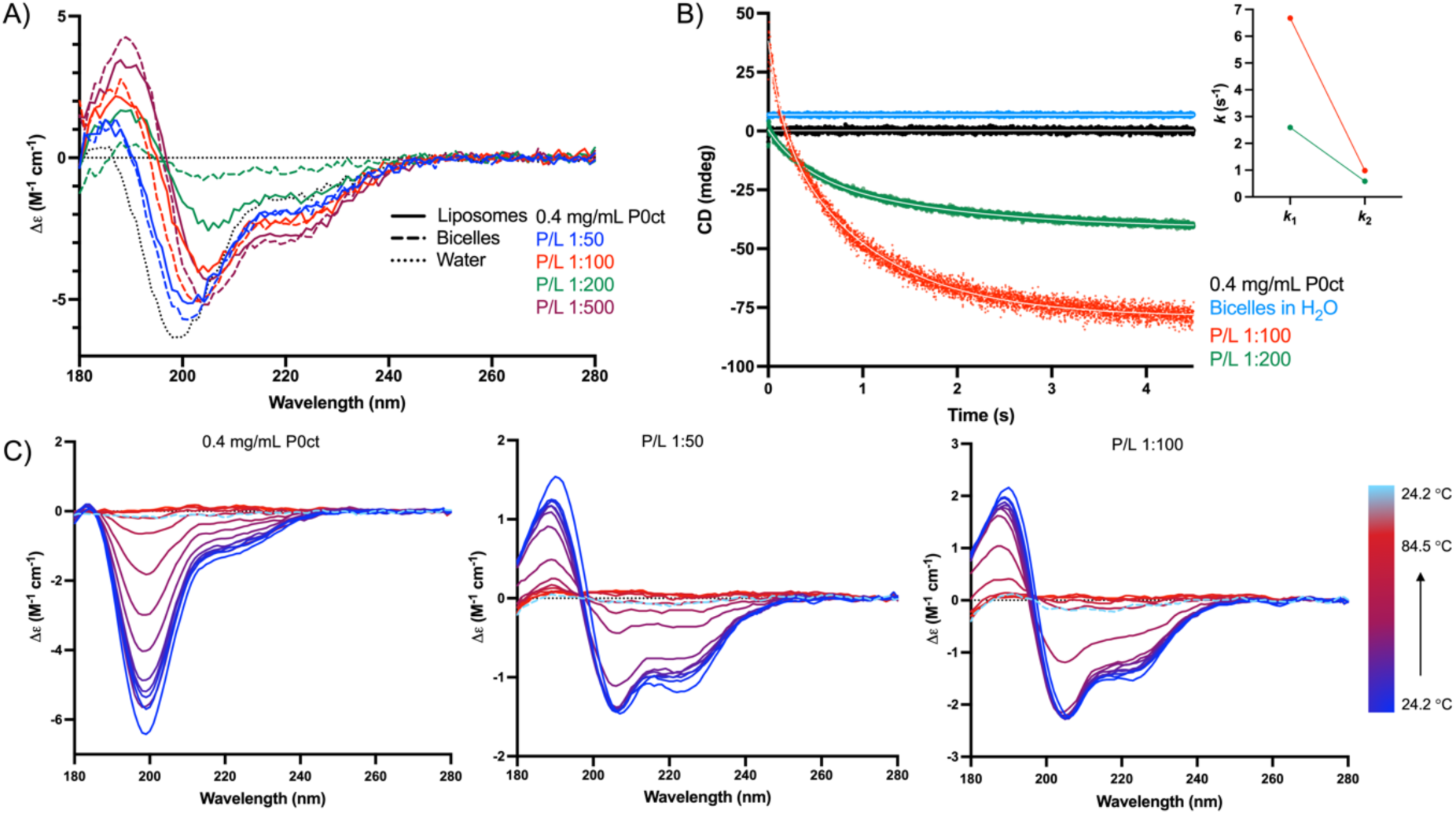
SRCD experiments. A) The folding of P0ct in the two models (DMPC:DMPG 1:1) was studied using SRCD with different P/L ratios. Additional spectra with various lipid mixtures are presented in Supplementary Figure 3. B) Stopped-flow kinetics of bicelle aggregation. The SRCD signal at 195-200 nm was monitored for 5 s using stopped-flow measurements. P0ct was measured in the absence and presence of bicelles at two protein-to-lipid ratios with 150 mM NaF. The two rate constants k1 and k2 for P/L 1:100 and P/L 1:200 are indicated in the inset. See Supplementary Table 1 for all rate constants and Supplementary Figure 2 for the Log spectra. C) Thermal measurements were executed on three samples with temperatures ranging from 24 – 84 °C; P0ct in solution (left), P/L 1:50 (middle), P/L 1:100 (right). Bicelles were used as the membrane model.

### Bicelles aggregate more slowly than liposomes

The kinetics of the initial P0ct-induced bicelle aggregation were measured with stopped-flow SRCD, as previously established (Raasakka et al., 2019a) (Figure 2B). When P0ct and bicelles were measured separately, the kinetics were flat, indicating no vesicle fusion or aggregation in the isolated samples, whereas when mixed at two different P/L ratios, a strong decay can be measured within a 5 s time frame. All samples were reproducible between shots and replicates with good overlay (see Supplementary Table 1 for two-phase decay parameters). The P0ct-bicelle sample data could be fitted using two rate constants: k_1_, describing a fast event (> 2 s^-1^), and k_2_, for a slower event (< 1 s^-1^) (Raasakka et al., 2019a). At P/L of 1:100, k_1_ is more than doubled compared to P/L of 1:200, while the slower rate constant is in a similar range (P/L 1:100 k_2_ = 0.98 s^-1^, P/L 1:200 k_2_ = 0.58 s^-1^). Hence, P0ct induces faster turbidity when less lipids are present. With liposomes, P/L ratio 1:200 has a k_1_ of 20.14 s^-1^ and a k_2_ of 1.12 s^-1^ (Raasakka et al., 2019a; Raasakka et al., 2019b). Why the differences in k_1_ between the two model systems are so substantial is not known, but the results suggest that the initial aggregation of liposomes happens at a faster rate than for the bicelles. This may be caused by larger interacting surface areas.

### P0ct keeps its folding at high temperatures

Thermal measurements on P2 have revealed that secondary structure was still observed after subjecting P2-bicelle aggregates to high temperatures (unpublished data); similarly, P2 remained folded in DMPC:DMPG liposomes up to +90 °C (Ruskamo et al., 2014). Without the lipid model present, P2 denatures completely at ∼+60 °C (Ruskamo et al., 2014). To see if P0ct behaves similarly, samples were subjected to thermal denaturation (Figure 2C) at 24 – 80 °C, followed by a final measurement after re-cooling back to 24 °C. In the absence of lipids, P0ct aggregates as the temperature rises, as evident from the decreasing CD signal intensity (apparent T_m_ 57.9 °C). In the bicelle-containing samples, secondary structure is prominent at 24 °C, and this helical structure is stable for several temperature increments before the signal disappears. High temperatures cause the protein-bicelle clusters to aggregate (apparent T_m_ 55.0 °C at P/L 1:50 and 61.4 °C at P/L 1:100). P0ct remains helical upon bicelle aggregation at higher temperatures, as the positions of the CD peaks do not change. Hence, melting temperatures from these experiments reflect sample aggregation rather than actual protein unfolding.

### High concentrations of P0ct fuse membranes instead of stacking

TEM was used to visualize the effects of P0ct on the different lipid models. At the lowest P/L ratios of either SUVs or bicelles, P0ct induces lipid aggregates differently in the two membrane-mimicking systems. With the highest concentration of P0ct (P/L 1:50), liposomes form into what resembles membrane sheets (Figure 3A, upper row), suggesting fusion of the free liposomes seen in control samples (Figure 1C). From several of the sheets, one can see “vesicle tails” protruding from the merged vesicles. This suggests that the protein is used up in the formation of the larger lipid structures, leaving some liposomes in an unfused state. At increasing lipid content, the sheets are no longer visible, but instead, clusters of vesicles can be seen. In the vesicle control sample, the vesicles are seen freely on the grid, while at P/L 1:100, some vesicles are still free in solution, but a larger amount is aggregated together in variable-sized assemblies. The same is true for P/L 1:200, although more free vesicles are observed.

**Figure 3:**
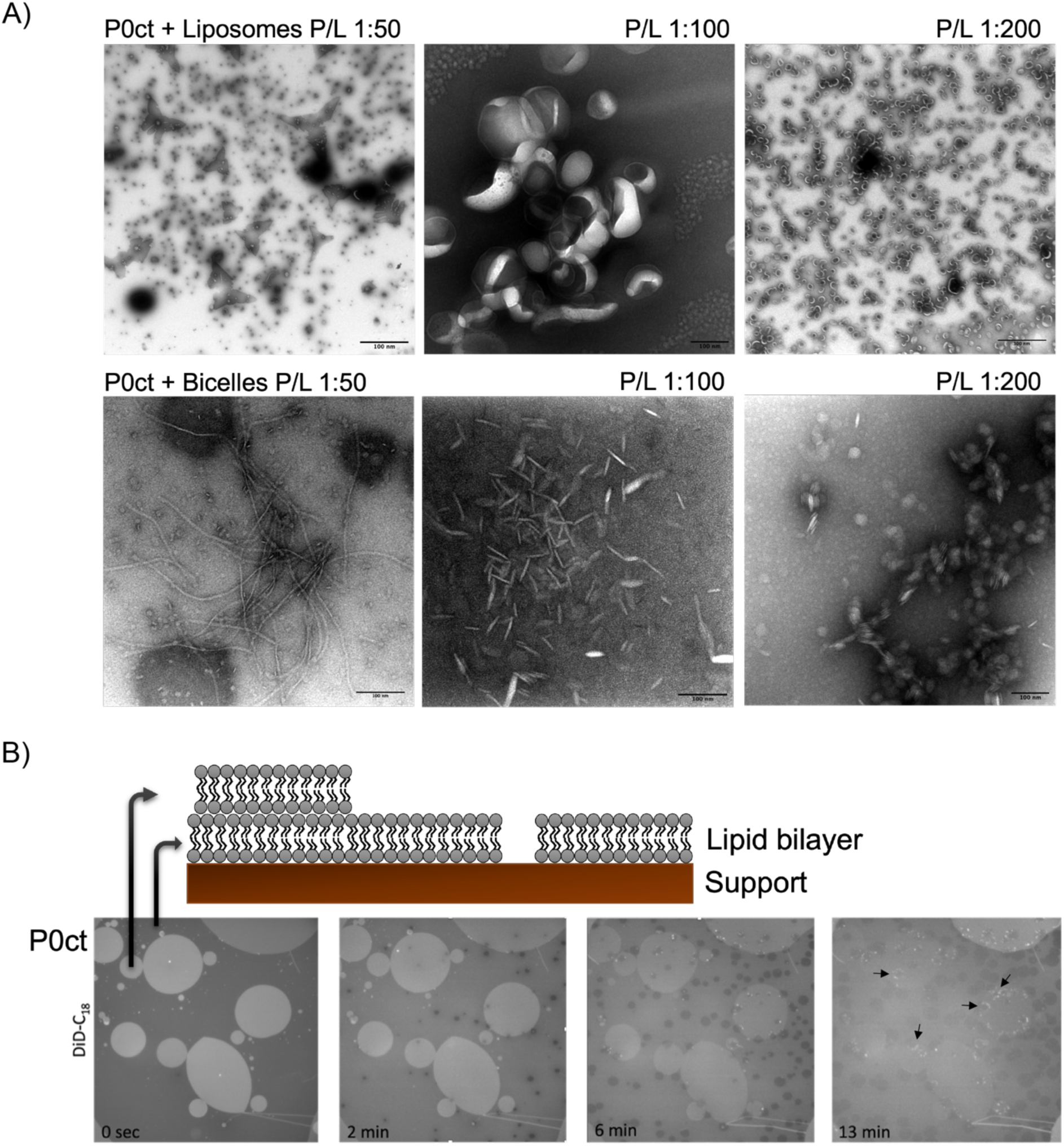
Effect of P0ct on membrane morphology. A) Liposomes on the top row, bicelles on the bottom row, at 3 different P/L ratios. P0ct fuses lipid particles differently depending on the membrane model and P/L ratio; P/L 1:50 assembles uniquely. With liposomes, P0ct induces sheath-like structures with protruding lipid “tail”, while the protein fuses bicelles together into filamentous threads. Control TEM images are in shown in Supplementary Figure 1. Scalebars are set to 100 nm. B) Fluorescence patch experiment. 10 μM of P0ct was added to DOPC:DOPS (9:1, 0.05% DiD-C18) and visualized for 15 min. Before adding protein (0 s), the patches and the lipid bilayer underneath are intact. The addition of P0ct induces cavities in the underlying lipid layers and accumulation of lipids in the developed gaps (bright spots). This formation was not observed in bilayer controls (not shown). Original movies of the experiment are shown in Supplementary Movies 1-2.

When using bicelles, the aggregates formed by P0ct appear different. In contrast to the membrane sheets, bicelles with the same concentration of P0ct (P/L 1:50) fuse into long filament-like structures (Figure 3A, lower row). Visually, there is a large difference in the appearance of the membrane-mimicking systems, when P0ct is added at high concentrations. The formation of filaments is lost when more lipids are present (P/L 1:100). The bicelles appear to be aggregated more randomly, and at P/L 1:200, stacks of bicelles can be observed.

For both lipid models, when the concentration of P0ct is high enough, either vesicles or bicelles merge, forming larger, relatively flat lipid structures. The membranes may be saturated with the protein in this case, which prevents their stacking. Similar conclusions were earlier drawn based on atomic force microscopy experiments (Raasakka et al., 2019c). For higher ratios of lipids, the appearance is more similar, more free lipids are seen when the lipid ratio increases. The results indicate that P0ct can affect lipid membrane morphology in a concentration-dependent manner. Since the above SRCD experiments indicated that at P/L 1:50, P0ct is less folded than at P/L 1:100, it appears that the membrane has reached saturation of its capacity to bind P0ct and induce its folding.

### P0ct alters membrane morphology

Time-lapse fluorescence microscopy was utilized to further investigate the protein-lipid interactions of P0ct on planar supported bilayers. Bilayers of DOPC and DOPS were prepared with a DOPC:DOPS ratio of 9:1, and 0.05% DiD-C18 probe was added to the lipid mixture as a fluorescent dye. Before the addition of P0ct, the supported bilayers stack on top of each other, forming double bilayer islands, without measurable malformations (Figure 3B, 0 s). The effects of P0ct addition were recorded for approximately 15 min. With time, the fluorescent bilayers become bleached by the prolonged light exposure. The first indication of P0ct influencing the bilayers appeared after ∼2 min. The first consequence of P0ct addition is the manifestation of holes/cavities in the primary bilayer (darkest shaded bilayer). These holes may be a sign of membrane condensation induced by P0ct, decreasing the area per lipid molecule. With time, more cavities are observed, and bright spots emerge at the edges of the holes (black arrows in Figure 3B), indicating the formation of new lipid structures. Over 15 min, the cavities grow, and more lipids accumulate at the edges. Bilayers without protein were filmed as control, but no changes in morphology were observed, implying that the changes in membrane structure are induced by P0ct. The lipid accumulation is only observed when there are secondary bilayers. In a similar experiment, myelin protein P2 induced the formation of new membrane structures (Uusitalo et al., 2021) without the generation of cavities; thus, the effects of these two PNS myelin MDL proteins on supported lipid bilayers are different.

### Evidence of hexagonal lipid crystals at high protein concentration

We performed SAXD with liposomes in previous studies on both P0ct and MBP (Raasakka et al., 2019b; Raasakka et al., 2017; Raasakka et al., 2019c), but did not previously employ bicelles. The first myelin protein-bicelle diffraction experiments were performed with P2 (Ruskamo et al., 2020), whereby a set of new diffraction peaks were observed upon the addition of P2 protein. The results were interpreted as the formation of semi-crystalline proteolipid structures. With liposomes, the typical peak induced by P0ct at around 70-75 Å is visible with its second-order peak, indicating ordered lamellar stacking (Figure 4A-B, black lines). The same repeat distance has been observed in previous work and is relatively consistent between measurements and protein concentrations (Raasakka et al., 2019b; Raasakka et al., 2019c). The 75 Å repeat distance corresponds to the thickness of cytoplasmic stacks in myelin (two cytoplasmic membrane leaflets and the protein phase in between) and is in the same range as reported for P2 and MBP (Campi et al., 2017; Raasakka et al., 2017; Ruskamo et al., 2020). For liposomes, the membrane thickness was measured to be 37.7 Å (Figure 1), and the lamellar periodicity was 73.8 Å and 75.3 Å for P/L 1:100 and P/L 1:50, respectively (Figure 4C). This denotes that the water gap, containing P0ct, between the bilayers is 36.2 Å and 38.2 Å for the two concentrations of P0ct; a higher concentration of P0ct increases the intermembrane distance. Changing the concentration of P0ct does not change the number of diffraction peaks seen in the experiment, but the narrowing of the peaks indicates a higher degree of order in the repeating units (Li et al., 2017). Liposomes have for a long time been the go-to model system in diffraction experiments, but using bicelles for these types of measurements is a good source of complementary information on how the proteins are arranged within a membrane system (Ruskamo et al., 2020).

**Figure 4:**
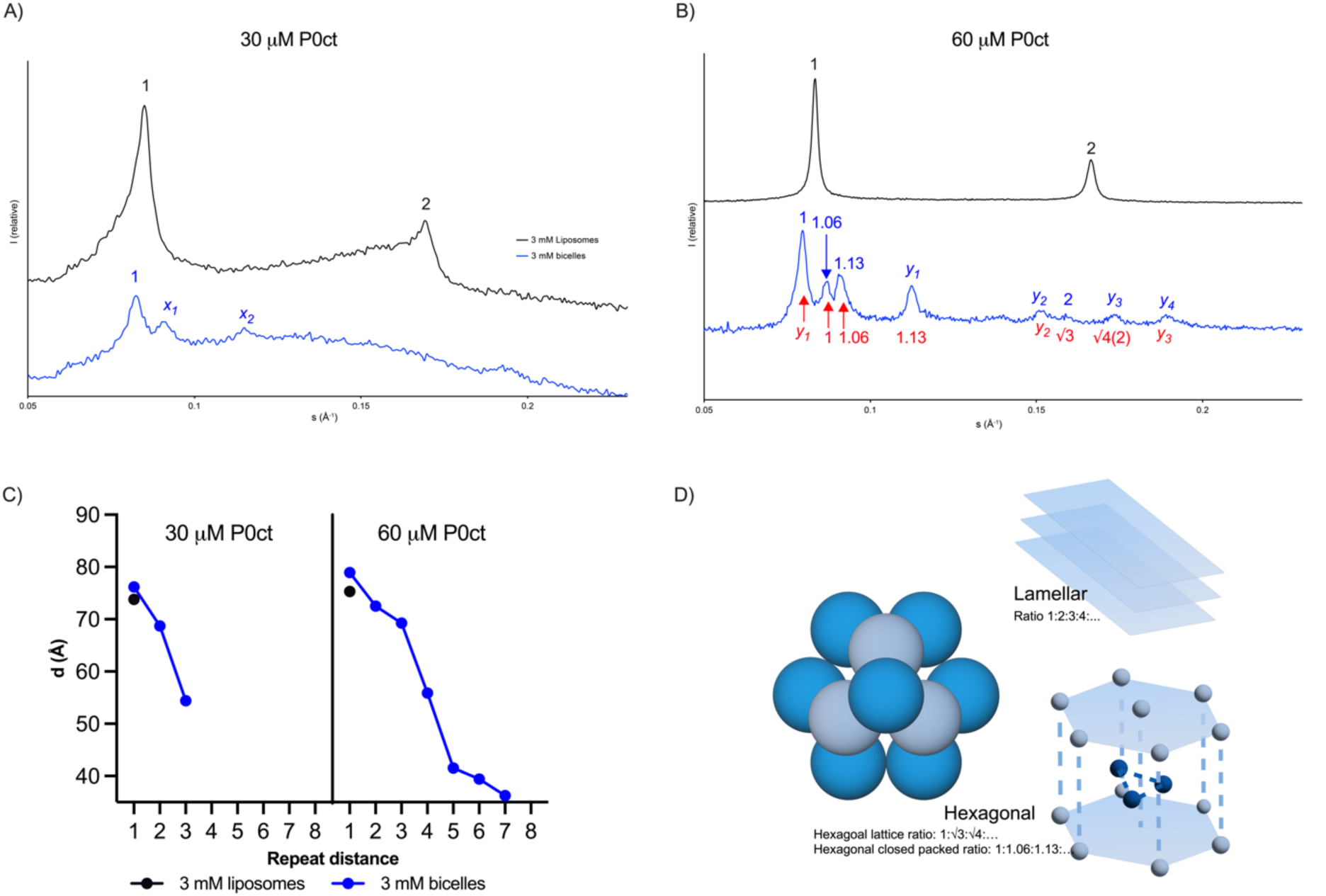
X-ray diffraction of P0ct with SUVs and bicelles. A-B) Adding 30 or 60 μM P0ct to liposomes (black lines) results in two major Bragg peaks, while the same concentration added to bicelles (blue lines) gives 3 or more Bragg peaks. Peak numbered with 1 indicates lamellar d-spacing. B) Blue and red labels for each peak reflect two potential approaches to analyzing the Bragg peaks. The spectra have been offset for clarity. C) The mean repeat distances, calculated from the s-values corresponding to the intensity summit of each Bragg peak, going from left to right corresponding to the peaks seen in A-B. One repeat distance is indicated for each liposome sample as the second peak is the second order and not a repeat distance by itself. D) Illustration of the main liquid crystal systems; lamellar ratio of 1:2:3:4:5… and the ratios of hexagonal lattice and hexagonal closed packed systems.

Up to eight peaks were observed for the highest concentration of P0ct when bicelles were used as the biomimetic analogue (Figure 4A-B, blue lines). No Bragg peaks were observed for P0ct alone or in complex with DPC detergent (Supplementary Figure 4). Three peaks appear at around 76.2, 68.7, and 54.4 Å when 30 µM of P0ct is used (P/L 1:100). In contrast, P/L 1:50 reveals eight peaks corresponding to 78.9, 72.5, 69.3, 55.9, 41.5, 39.4, 36.3, and 33.2 Å. The lamellar d-spacing of P/L 1:100 is peak 1 with a d-spacing of 76.2 Å, while the two additional peaks, labelled x_1_ (68.70 Å) and x_2_ (54.40 Å), did not fit a lamellar peak ratio of 1:2:3:4. The lamellar repeat distance in the bicelle system was therefore ∼3 Å larger than in the liposome system. The mean membrane thickness of the bicelles (*d_m,MKP_)* was estimated to be 34.8 Å; thus, the water gap was 41.4 Å for P/L 1:100, an increase of 5.2 Å compared to the liposome sample (an increase of 6.0 Å for P/L 1:50). The peak pattern for P/L 1:50 corresponds best to the presence of a hexagonal lattice system (Figure 4D). Two positions of the first-order lamellar peak 1 (blue and red labels in Figure 4B) reveal several peak ratios matching a hexagonal system, potentially mixed with a lamellar phase (Figure 4B, blue and red respectively). Regardless of the peak 1 position, there are peaks not fitting to any lattice system tested, marked with y_1-4_. For both bicelle samples, the unidentified peaks (x_1-2_, y_1-4_) might be evidence that unidentified non-lamellar phases occur within the bicelle cluster, collectively with the lamellar d-spacing and the hexagonal lattice system. Taken together, when mixed with liposomes, P0ct induces lamellar stacking of the membranes, while in the bicelle system, a mixture of lipid phases is observed.

### Purification and initial characterisation of recombinant human full-length P0

The cytoplasmic tail of P0 has been extensively studied, while hP0 has not been purified for structural studies before. P0ct can be produced recombinantly in *E. coli*, whereas hP0 is a greater challenge, as membrane proteins are in general difficult to purify and require detergent to remain in solution (Lin & Guidotti, 2009). We expressed and purified hP0 using insect cells and a cleavable GFP tag. DDM was found to be a suitable detergent to obtain pure protein. In short, the membrane homogenate was incubated with DDM to solubilize hP0, and IMAC was executed before and after cleavage of the tag, before a final SEC step to obtain pure protein. Mass spectrometry was executed on the final product to verify the correct protein.

The Ig-like domain of P0 has been crystallized and characterized at high resolution (Liu et al., 2012; Shapiro et al., 1996), while biophysical characterization has been done for P0ct (Krokengen et al., 2023; Luo et al., 2007; Raasakka et al., 2019a; Raasakka et al., 2019b; Raasakka et al., 2019c). Consequently, this key protein of PNS myelin has only been studied in parts, and no experimental information on full-length P0 exists, apart from low-resolution electron cryomicrographs of liposome-reconstituted P0 purified from bovine nerves and reconstituted into liposomes (Raasakka et al., 2019c). Since more experiments are required to fully understand P0 function in myelin, we present the first structural characterization of recombinant hP0 (Figure 5). SEC-SAXS data were acquired for hP0 in DDM micelles; SEC facilitated the separation of empty micelles from protein-detergent complexes. SAXS data for hP0 can be seen in Figure 5A, and from the normalized Kratky plot, hP0 appears to be folded with flexible domains (Figure 5B). R_g_ derived from the SAXS data was 38.83 ± 0.7 Å, with a D_max_ of 119.10 nm (Figure 5C). To obtain a model for the protein-detergent complex based on the SAXS data, several modelling programs were tested; a hybrid model generated in CORAL (Petoukhov et al., 2012) had the best fit (Figure 5A,D). An AlphaFold2 (Jumper et al., 2021) model for hP0 was generated as well as a 3D reconstruction of the electron density from the experimental SAXS data. SEC-SAXS data were also acquired for highly concentrated, pure P0ct (Supplementary Figure 5) to supplement the findings; an AlphaFold2 model of P0ct has been discussed before (Krokengen et al., 2023). P0ct was elongated and flexible, showing a mixture of more extended and compacted conformations (Supplementary Figure 5).

**Figure 5:**
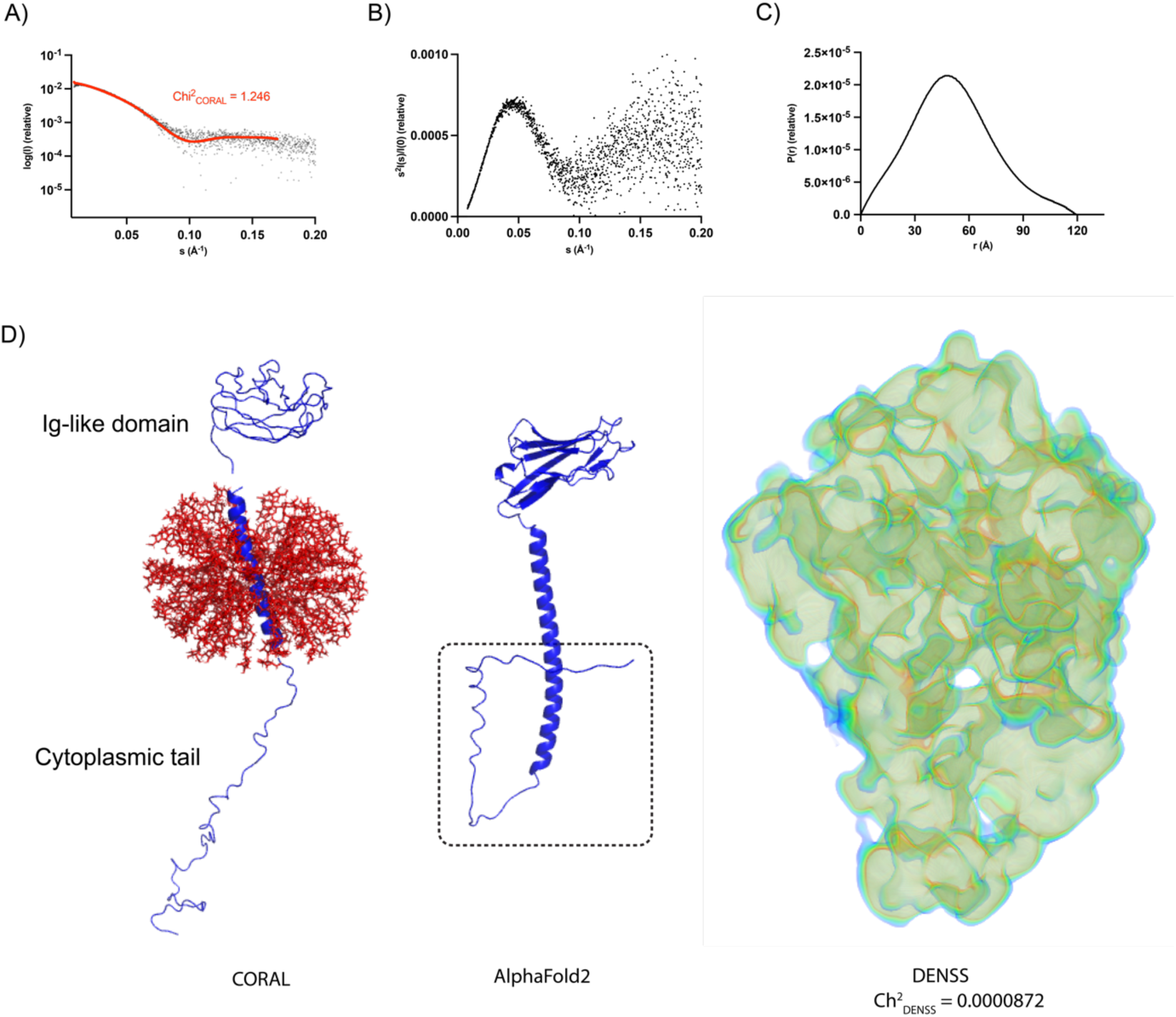
Characterisation of hP0 in DDM. A) SAXS curve and the CORAL fit, B) Kratky plot, and C) distance distribution. D) Hybrid CORAL model of hP0 with DDM molecules (left), AlphaFold2 model of hP0 given only the sequence as input (UniProt ID: P25189) (Jumper et al., 2021) (N-terminal region removed for clarity, cytoplasmic tail encircled) (middle), and a 3D reconstruction of the electron density profile (right).

In the SAXS-based model of the hP0-DDM complex (Figure 5D), a detergent belt surrounds the TM helix, while the Ig-like domain and the cytoplasmic tail are placed on opposite sides of the micelle. The SAXS-based model is similar, but not identical, to the AlphaFold2 model. At the C-terminal end, the cytoplasmic tail is modelled as elongated based on SAXS; in the AlphaFold2 model, it turns back towards the TM helix. The AlphaFold2 per-residue confidence score (pLDDT) and the predicted aligned error (PAE) are shown in Supplementary Figure 6A. Molecular dynamics simulations (Eastman et al., 2017; Tesei & Lindorff-Larsen, 2023; Tesei et al., 2024) of hP0 similarly conclude with an R_g_ of 42.8 Å. A DENSS electron density fit of the SEC-SAXS data is shown in Figure 5D (fitting of the DENSS map is shown in Supplementary Figure 6B), indicating an increased volume, showing the DDM micelle around the TM helix. To conclude, modelling of full-length hP0 based on both predictions and SAXS data indicates a folded extracellular domain, a TM helix in a detergent micelle, and a flexible cytoplasmic tail. The dimensions of P0ct in solution indicate that it must obtain a more compact structure, when bound between two membranes at the myelin MDL.

## Discussion

Accurate ensheathment of axons depends on several proteins, whereby the P0 glycoprotein is vital for myelination and myelin compaction in the PNS (Kister & Kister, 2023; Prada et al., 2012; Raasakka & Kursula, 2020). As an integral membrane protein, P0 functions as an anchoring protein on both sides of the membrane; it adheres extracellular membrane leaflets together through oligomerization of the Ig-like domains, and on the opposite side, P0ct partially folds and inserts into the lipid membrane (Eichberg, 2002; Gao et al., 2000; Plotkowski et al., 2007; Wong & Filbin, 1994). Nonetheless, the P0 molecular mechanism remains a mystery. We previously investigated P2 using bicelles as a membrane model (Ruskamo et al., 2020) to yield new evidence of myelin structural assembly. Here, we report on the use of different membrane mimics in the investigation of P0ct function within the MDL of PNS myelin. We also present a preliminary structural analysis of full-length human P0.

### Membrane morphology and P0

Although liposomes and bicelles differ mostly in shape, it is evident that the model system used has a notable impact on P0ct-lipid interactions. Changing the membrane model does not influence the secondary structure of P0ct, but as visualized in TEM (Figure 3A), a high concentration of P0ct induces the formation of either sheet-or filament-like structures. Filament structures have been observed with liposomes at higher P/L ratios (Raasakka et al., 2019c) but were only observed with bicelles here. The thickness of the filaments was 10-20 nm, matching earlier findings (Raasakka et al., 2019c). With either model system, high concentrations of P0ct appear to cause membrane fusion instead of membrane aggregation, which suggests the importance of correct protein concentration for myelin membrane wrapping and stacking. Linked to this observation, at P/L 1:50, P0ct was less folded thatn at higher lipid concentrations, suggesting a relationship on the P0ct folding state, its concentration on the membrane, and its effect ofn membrane structure.

Using liposomes, the Bragg peaks indicated the presence of a lamellar structure, with a more ordered structure with the addition of more P0ct (Figure 4A-B) (Raasakka et al., 2019b; Raasakka et al., 2019c). However, for bicelles, more peaks appear at higher protein concentrations, which cannot be explained with simple lamellar d-spacing (Figure 4). Amphiphilic lipids can have polymorphism for other phases including hexagonal, bicontinuous cubic, sponge, and discrete micellar phases (Kulkarni et al., 2011; Lombardo et al., 2015), also in complex with other molecules (Bilalov et al., 2012). When comparing various lattice systems, it became apparent that P0ct modulates lipid phase transitions depending on which model system is used, *i.e.* liposomes or bicelles. In the 1980s, Kruijff and Cullis found that both the mitochondrial protein cytochrome C and the attachment factor poly-L-lysine caused hexagonal phase transition in membranes containing certain negatively charged lipids (de Kruijff & Cullis, 1980a, 1980b). Following either blue or red labels in Figure 4B, the hexagonal lattice system matches 2-4 peaks induced by P0ct, while two Bragg peaks fit a lamellar lattice (Chen et al., 2019; Hsu et al., 2020). The Ig-like domain of P0 is understood to stack membranes in a zipper-like fashion (Raasakka & Kursula, 2020; Raasakka et al., 2019c), but the P0ct mode of binding is not fully known. The basic peptide poly-L-lysine and MBP are both thought to not only affect the membrane surface but also to interlink with the hydrophobic tails of the lipids (Boggs et al., 1981; Carrier & Pézolet, 1984), increasing the mobility of the lipid alkyl chains (Natali et al., 2002). Binding of P0ct leads to thermotropic modifications of lipid membranes (Raasakka et al., 2019b; Raasakka et al., 2019c); in addition, structural alterations appear in bicelle membranes visualized by TEM and confirmed by SAXD. DPC does not cause diffraction peaks on its own (Supplementary Figure 4), so the lipid phase transition is due to protein-lipid interactions in the model system. Many peaks cannot be explained by simple lamellar or hexagonal lattices; thus, other systems need to be involved as well. There is evidence that cubic phases can be established close to the L-H_II_ phase transition (Rappolt et al., 2003), and neutron reflectometry revealed that P0ct inserts into supported DMPC:DMPG lipid bilayers (Raasakka et al., 2019b). Consequently, it is feasible that P0ct embeds part of itself close to the core of the bicelle generating the increased movement needed for the phase transition towards hexagonal and other connected lattice systems. Another potential explanation is that the unknown peaks can be attributed to the protein organization within the lipid membranes. Diffraction peaks have been attributed to the organization of MBP bound to multilamellar vesicles (Shaharabani et al., 2016), and the same can be true for P0ct. Further investigation, using complementary techniques, will be needed to reveal the full mechanism.

Induced hexagonal phase transition has not been studied for P0, but it can be a feature of a disrupted myelin structure potentially relevant for neuropathies associated with P0, as found with other myelin proteins, including MBP, PMP22, and PLP. Shaharabani *et al*. found that MBP, a major protein in CNS and associated with multiple sclerosis, induced hexagonal phase transition in a disease-like lipid membrane composition (Shaharabani et al., 2016). The presence of the hexagonal phase reduced with lowered MBP concentration, indicating a relationship between them; nevertheless, temperature, lipid composition, and salt concentration also affected lipid organization (Shaharabani et al., 2018). Like for MBP in either lipid composition, native or disease-like (Shaharabani et al., 2016), the addition of P0ct reveals several correlation peaks demonstrating enhanced organization of the lipid membranes. Wrabetz *et al*. studied peripheral nerve myelination in mice concerning the expression of P0. The results revealed a dose-dependent dysmyelinating neuropathy with a threshold for dysmyelination between 30-80% overexpression of P0, indicating that normal myelinogenesis requires a strictly regulated expression of P0 (Wrabetz et al., 2000). This observation may be linked to the different membrane morphologies observed at high concentrations of P0ct here. Dysmyelination has been observed when overexpressing PLP, causing Pelizaeus-Merzbacher disease (Nave & Boespflug-Tanguy, 1996), and PMP22 overexpression is linked to CMT1A (Robaglia-Schlupp et al., 2002). Although most neuropathies related to P0 involve mutations found in the Ig-like domain or P0ct (Avila et al., 2010; Bai et al., 2018; Blanquet-Grossard et al., 1996; Callegari et al., 2019; Fabrizi et al., 2006; Filbin et al., 1999; Planté-Bordeneuve et al., 2001; Prada et al., 2012; Raasakka et al., 2019b; Schneider-Gold et al., 2010), rare cases of increased P0 gene dosage have been reported, in which patients appeared with motor and sensory polyneuropathy (Chen et al., 1994; Maeda et al., 2012). A concentration-and folding-dependent effect on lipid membrane properties could, therefore, be studied further as a molecular disease mechanism of de-and dysmyelination.

Having observed the filament-like membrane structures in the TEM images, we propose a potential mechanism for the proteolipid structure (Figure 6). When P0ct is added to either liposomes or bicelles at lower concentrations, P0ct binds to the membrane and induces lamellar stacking. With high concentrations, when the membrane surface starts to get saturated with protein, the bicelles start to fuse. Simultaneously with the folding of P0ct, the bicelles fuse, establishing longer filaments where “pockets” within the filament contain hexagonal phases, as the Bragg peaks reflect. The filaments do not seem to adhere to each other, which further supports the notion that the protein is enclosed by lipids, as postulated for cytochrome C (de Kruijff & Cullis, 1980a).

**Figure 6:**
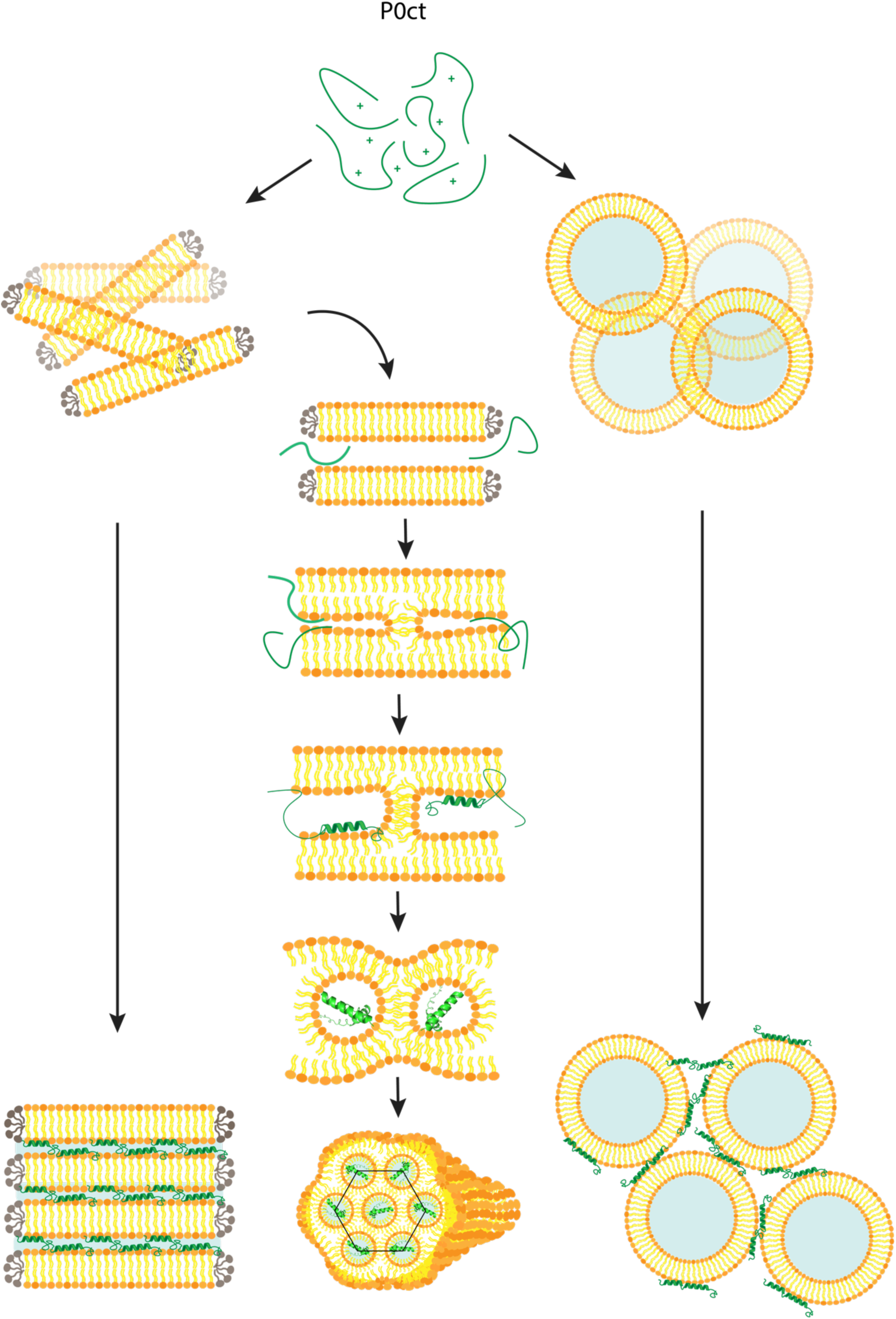
Schematic illustration of the potential hexagonal folding of P0ct-bicelle samples. When P0ct is added to bicelles, we propose a potential mechanism for a hexagonal filament induced by P0ct. At lower P/L ratios, most diffraction peaks are consistent with a lamellar lattice for either bicelles or liposomes (far right and left side). At higher protein concentrations, P0ct fuses bicelles, forming longer filaments where a network of “pockets” inside harbours P0ct (middle).

In myelin, P0 is mainly thought of as an adhesion molecule, but based on TEM and the patch experiments in combination with X-ray diffraction, one can speculate that P0 might be an important factor in the migration of Schwann cell membrane and its elongation along the axon, as postulated by Wrabetz *et al*. (Wrabetz et al., 2000). Not only does P0ct induce a membrane fusion at low P/L ratios, but it also affects membrane properties (Figure 3B). Annexin A1 and A2 have been reported to induce the formation of double-membrane structures (Berg Klenow et al., 2021; Boye et al., 2018), which was also shown for myelin protein P2 (Uusitalo et al., 2021). At lower concentrations of P0ct, holes and lipid accumulations were seen, while at the same concentration of P2 (unpublished data), the formation of new membrane structures can be observed, indicating myelin protein concentration-dependent changes in membrane morphology. The possible membrane-condensing effect observed for P0ct on supported planar membranes, together with its effect on membrane phase transition behaviour observed before (Raasakka et al., 2019c), may be related to molecular mechanisms of myelin formation.

### P0 structure starts to unfold

Even though P0 is recognized as a core building block of PNS myelin, full-length human P0 has not been recombinantly purified before, and a high-resolution structure of hP0 is not available. As a first step towards structural studies on hP0 and its complexes, we reported here the expression, purification, and initial SEC-SAXS characterisation of hP0. A Kratky plot of the SAXS curve shows a folded protein with some flexible domains. hP0 consists of an N-terminal signal sequence (residue 1-29) and three domains; the Ig-like domain (residue 30-153), the TM helix (residue 154-179) and the cytoplasmic tail (residue 180-248); the Ig-like domain is the only one crystallized (rat: PDB ID 1NEU, human: PDB ID 3OAI) (Liu et al., 2012; Shapiro et al., 1996). Crystallization of the extracellular domain revealed an Ig-like domain comprising ten β-strands arranged as two β-sheets displaying a hydrophobic core featured by the immunoglobulin family (Halaby & Mornon, 1998; Liu et al., 2012; Shapiro et al., 1996). In the SAXS-based model, a monomeric fraction of hP0 was analysed, and hence, no oligomerisation of the extracellular domain was observed.

The arrangement of the P0 Ig-like domains in the IPL of PNS myelin has been an open question. In our earlier work, cryo-EM revealed a zipper-like arrangement of the Ig-like domains (Raasakka et al., 2019c), which challenged existing models of the IPL. The conserved residues Arg74 and His81 of the Ig-like domains interact, causing the adhesion needed for a tight compartment (Inouye & Kirschner, 2016; Raasakka & Kursula, 2020). Mutations linked to these amino acids, together with several others in the same domain, are linked to reduced P0 function in several types of CMT (Callegari et al., 2019; Filbin et al., 1999; Grandis et al., 2008; Hattori et al., 2003; Prada et al., 2012; Sorour et al., 1997).

The last residues at the end of the extracellular domain are disordered and believed to supply flexibility with respect to the membrane surface (Shapiro et al., 1996). 26 residues make up the P0 TM helix, engulfed by a DDM micelle in the model. The helix has a glycine zipper motif (GxxxGxxxG) commonly found in membrane-associated proteins (Kim et al., 2005; Plotkowski et al., 2007). The TM helix undergoes dimerization, which supports oligomerisation of the Ig-like domains on the same membrane (Plotkowski et al., 2007). Two mutations associated with CMT (G*134*R) and Dejerine-Sottas syndrome (G*138*R) reside within the TM domain, having substantial effects on P0 membrane insertion and dimerization (Plotkowski et al., 2007).

AlphaFold2 starts the TM helix at the correct residue (Tyr154) but models the TM domain and the cytoplasmic domain as one elongated helix with a short disordered tail. A part of the long helix in the AlphaFold2 model (Figure 5D, black dotted square) belongs to P0ct. In fact, an amphipathic helix is believed to be separated from the TM helix by a small hinge necessary for its flexibility to bind alongside the membrane surface (D’Urso et al., 1990; Krokengen et al., 2023). When P0ct is given as a single sequence, AlphaFold2 models P0ct to contain helical segments (Krokengen et al., 2023), which has been observed experimentally when bound to lipid membranes (Raasakka et al., 2019a; Raasakka et al., 2019b; Raasakka et al., 2019c). How P0ct binds and resides within the MDL of myelin is still not fully known. One important detail from the hybrid model is that the cytoplasmic tail is extended and not bound to the DDM micelle and appears to have the flexibility to bind to a second membrane. If the P0ct folded upon the same micelle, it would be seen with a smaller maximum dimension (D_max_), with dimensions similar to the AlphaFold2 model. The 3D electron density profile modelled with DENSS supports the notion of an elongated profile of the cytoplasmic tail protruding down from the helix-DDM assembly.

### Concluding remarks

P0 is adhesive on both sides of the PNS myelin membrane; a zipper-like assembly of the extracellular domains is accompanied by the intrinsically disordered P0ct in a tight space between two cytoplasmic membrane leaflets. We have provided new information on the interactions of P0ct with lipids, revealing concentration-dependent effects on membrane morphology and lipid phase behaviour. Such phenomena may be relevant when the myelin membrane grows and wraps around the axon, as well as when the multilayered membrane compacts to produce mature myelin. The presence of human disease-linked mutations in the P0ct domain highlights the importance of the interactions of this small, flexible segment with lipid membranes. Now that we can purify full-length human P0, these findings can be explored in a more native-like setting needed for revealing molecular mechanisms in myelinogenesis and myelin disease.

## Acknowledgements

We acknowledge the use of the Core Facility for Biophysics, Structural Biology, and Screening (BiSS) at the University of Bergen, which has received infrastructure funding from the Research Council of Norway (RCN) through NORCRYST (grant number 245828) and NOR–OPENSCREEN (grant number 245922). In addition, we are grateful to the Molecular Imaging Center (MIC) at the Department of Biomedicine, University of Bergen, for providing access to instrumentation. We acknowledge MAX IV Laboratory for time on the Beamline COSAXS under Proposal (ID 20220497/20200757). Research conducted at MAX IV, a Swedish national user facility, is supported by the Swedish Research Council under contract 2018-07152, the Swedish Governmental Agency for Innovation Systems under contract 2018-04969, and Formas under contract 2019-02496. We gratefully acknowledge the synchrotron radiation facilities and the beamline support at ASTRID2. Parts of the experiments carried out at ASTRID2 were under the support of the EU H2020 MOSBRI research and innovation program This project has received funding from the European Union’s Horizon 2020 research and innovation programme under grant agreement No 101004806, MOSBRI transnational access proposal ID MOSBRI-2022-137. The assistance of Nykola C. Jones and Søren Vrønning Hoffman at the ASTRID2 synchrotron (Aarhus, Denmark) is highly appreciated. The authors would like to thank Diamond Light Source beamline B21 and principal beamline scientist Nathan Cowieson. We acknowledge financial support from the University of Bergen (Norway) and several travel grants provided by BioCat, The Norwegian national graduate school in Biocatalysis. The BIOPROM project is funded by the Research Council of Norway (project number 324877).

## Supplementary information

Supplementary Figure 1: Control TEM images

Supplementary Figure 2: SRCD control experiments and time scale of P0ct-induced initial lipid turbidity

Supplementary Figure 3: SRCD measurements of several protein-to-lipid ratios of P0ct in various DMPC:DMPG lipid mixtures with decreasing ratios of negatively charged lipids

Supplementary Figure 4: SAXD measurements of various protein-to-lipid ratios of P0ct and 3 mM DPC detergent.

Supplementary Figure 5: SEC-SAXS measurements and model of 14.4 mg/mL P0ct.

Supplementary Figure 6: AlphaFold2 model and DENSS curve fit of full-length P0 (hP0)

Supplementary Table 1: Non-linear fit parameters

Supplementary Table 2: SAXS parameters for P0ct and full-length P0 Supplementary Movie 1: patch experiment of 10 μM P0ct (as seen in Figure 3B)

Supplementary Movie 2: control experiments of patches

## References

Akbarzadeh, A., Rezaei-Sadabady, R., Davaran, S., Joo, S. W., Zarghami, N., Hanifehpour, Y., Samiei, M., Kouhi, M., & Nejati-Koshki, K. (2013). Liposome: classification, preparation, and applications. Nanoscale Res Lett, 8(1), 102. 10.1186/1556-276x-8-102

Avila, R. L., D’Antonio, M., Bachi, A., Inouye, H., Feltri, M. L., Wrabetz, L., & Kirschner, D. A. (2010). P0 (protein zero) mutation S34C underlies instability of internodal myelin in S63C mice. J Biol Chem, 285(53), 42001–42012. 10.1074/jbc.M110.166967

Bai, Y., Wu, X., Brennan, K. M., Wang, D. S., D’Antonio, M., Moran, J., Svaren, J., & Shy, M. E. (2018). Myelin protein zero mutations and the unfolded protein response in Charcot Marie Tooth disease type 1B. Ann Clin Transl Neurol, 5(4), 445–455. 10.1002/acn3.543

Beaugrand, M., Arnold, A. A., Hénin, J., Warschawski, D. E., Williamson, P. T. F., & Marcotte, I. (2014). Lipid Concentration and Molar Ratio Boundaries for the Use of Isotropic Bicelles. Langmuir, 30(21), 6162–6170. 10.1021/la5004353

Berg Klenow, M., Iversen, C., Wendelboe Lund, F., Mularski, A., Busk Heitmann, A. S., Dias, C., Nylandsted, J., & Simonsen, A. C. (2021). Annexins A1 and A2 Accumulate and Are Immobilized at Cross-Linked Membrane-Membrane Interfaces. Biochemistry, 60(16), 1248–1259. 10.1021/acs.biochem.1c00126

Bharadwaj, M., & Bizzozero, O. A. (1995). Myelin P0 Glycoprotein and a Synthetic Peptide Containing the Palmitoylation Site Are Both Autoacylated. Journal of Neurochemistry, 65(4), 1805–1815. 10.1046/j.1471-4159.1995.65041805.x

Bilalov, A., Olsson, U., & Lindman, B. (2012). Complexation between DNA and surfactants and lipids: phase behavior and molecular organization [10.1039/C2SM26553B]. Soft Matter, 8(43), 11022–11033. 10.1039/C2SM26553B

Bizzozero, O. A., Fridal, K., & Pastuszyn, A. (1994). Identification of the Palmitoylation Site in Rat Myelin P0 Glycoprotein. Journal of Neurochemistry, 62(3), 1163–1171. 10.1046/j.1471-4159.1994.62031163.x

Blanquet-Grossard, F., Pham-Dinh, D., Dautigny, A., Latour, P., Bonnebouche, C., Diraison, P., Chapon, F., Chazot, G., & Vandenberghe, A. (1996). Charcot-Marie-Tooth type 1B neuropathy: a mutation at the single glycosylation site in the major peripheral myelin glycoprotein Po. Hum Mutat, 8(2), 185–186. 10.1002/(sici)1098-1004(1996)8:2<185::Aid-humu13>3.0.Co;2-z

Boggs, J. M., Stamp, D., & Moscarello, M. A. (1981). Interaction of myelin basic protein with dipalmitoylphosphatidylglycerol: dependence on the lipid phase and investigation of a metastable state. Biochemistry, 20(21), 6066–6072. 10.1021/bi00524a023

Borrell, J. H., Montero, M. T., Morros, A., & Domènech, Ò. (2015). Unspecific membrane protein–lipid recognition: combination of AFM imaging, force spectroscopy, DSC and FRET measurements. Journal of Molecular Recognition, 28(11), 679–686. 10.1002/jmr.2483

Boye, T. L., Jeppesen, J. C., Maeda, K., Pezeshkian, W., Solovyeva, V., Nylandsted, J., & Simonsen, A. C. (2018). Annexins induce curvature on free-edge membranes displaying distinct morphologies. Scientific Reports, 8(1), 10309. 10.1038/s41598-018-28481-z

Brooks, B. R., Brooks, C. L., 3rd, Mackerell, A. D., Jr., Nilsson, L., Petrella, R. J., Roux, B., Won, Y., Archontis, G., Bartels, C., Boresch, S., Caflisch, A., Caves, L., Cui, Q., Dinner, A. R., Feig, M., Fischer, S., Gao, J., Hodoscek, M., Im, W., … Karplus, M. (2009). CHARMM: the biomolecular simulation program. J Comput Chem, 30(10), 1545–1614. 10.1002/jcc.21287

Callegari, I., Gemelli, C., Geroldi, A., Veneri, F., Mandich, P., D’Antonio, M., Pareyson, D., Shy, M. E., Schenone, A., Prada, V., & Grandis, M. (2019). Mutation update for myelin protein zero-related neuropathies and the increasing role of variants causing a late-onset phenotype. J Neurol, 266(11), 2629–2645. 10.1007/s00415-019-09453-3

Campi, G., Gioacchino, M. D., Poccia, N., Ricci, A. M. V., Burghammer, M., & Bianconi, A. (2017). Intrinsic dynamical fluctuations of PNS myelin.

Carrier, D., & Pézolet, M. (1984). Raman spectroscopic study of the interaction of poly-L-lysine with dipalmitoylphosphatidylglycerol bilayers. Biophysical journal, 46(4), 497–506. 10.1016/s0006-3495(84)84047-3

Chan, Y. H., & Boxer, S. G. (2007). Model membrane systems and their applications. Curr Opin Chem Biol, 11(6), 581–587. 10.1016/j.cbpa.2007.09.020

Chen, H., Kusyk, C. J., Tuck-Muller, C. M., Martinez, J. E., Dorand, R. D., & Wertelecki, W. (1994). Confirmation of proximal 1q duplication using fluorescence in situ hybridization. American Journal of Medical Genetics, 50(1), 28–31. 10.1002/ajmg.1320500106

Chen, L.-T., Chen, C.-Y., & Chen, H.-L. (2019). FCC or HCP: The stable close-packed lattice of crystallographically equivalent spherical micelles in block copolymer/homopolymer blend. Polymer, 169, 131–137. 10.1016/j.polymer.2019.02.041

Choi, B. O., Lee, M. S., Shin, S. H., Hwang, J. H., Choi, K. G., Kim, W. K., Sunwoo, I. N., Kim, N. K., & Chung, K. W. (2004). Mutational analysis of PMP22, MPZ, GJB1, EGR2 and NEFL in Korean Charcot-Marie-Tooth neuropathy patients. Hum Mutat, 24(2), 185–186. 10.1002/humu.9261

Conn, C. E., de Campo, L., Whitten, A. E., Garvey, C. J., Krause-Heuer, A. M., & van ’t Hag, L. (2020). Membrane Protein Structures in Lipid Bilayers; Small-Angle Neutron Scattering With Contrast-Matched Bicontinuous Cubic Phases. Front Chem, 8, 619470. 10.3389/fchem.2020.619470

Corin, K., & Bowie, J. U. (2020). How bilayer properties influence membrane protein folding. Protein Science, 29(12), 2348–2362. 10.1002/pro.3973

Corin, K., & Bowie, J. U. (2022). How physical forces drive the process of helical membrane protein folding. EMBO reports, 23(3), e53025. 10.15252/embr.202153025

Courreges, C., Nallet, F., Dufourc, E. J., & Oda, R. (2011). Unprecedented Observation of Days-Long Remnant Orientation of Phospholipid Bicelles: A Small-Angle X-ray Scattering and Theoretical Study. Langmuir, 27(15), pp. 9122–9130. 10.1021/la1050817

Cowieson, N. P., Edwards-Gayle, C. J. C., Inoue, K., Khunti, N. S., Doutch, J., Williams, E., Daniels, S., Preece, G., Krumpa, N. A., Sutter, J. P., Tully, M. D., Terrill, N. J., & Rambo, R. P. (2020). Beamline B21: high-throughput small-angle X-ray scattering at Diamond Light Source. Journal of Synchrotron Radiation, 27, 1438–1446. 10.1107/S1600577520009960

D’Urso, D., Brophy, P. J., Staugaitis, S. M., Gillespie, C. S., Frey, A. B., Stempak, J. G., & Colman, D. R. (1990). Protein zero of peripheral nerve myelin: biosynthesis, membrane insertion, and evidence for homotypic interaction. Neuron, 4(3), 449–460. 10.1016/0896-6273(90)90057-m

D’Urso, D., Ehrhardt, P., & Müller, H. W. (1999). Peripheral myelin protein 22 and protein zero: a novel association in peripheral nervous system myelin. J Neurosci, 19(9), 3396–3403. 10.1523/jneurosci.19-09-03396.1999

de Kruijff, B., & Cullis, P. R. (1980a). Cytochrome c specifically induces non-bilayer structures in cardiolipin-containing model membranes. Biochim Biophys Acta, 602(3), 477–490. 10.1016/0005-2736(80)90327-2

de Kruijff, B., & Cullis, P. R. (1980b). The influence of poly(l-lysine) on phospholipid polymorphism Evidence that electrostatic polypeptide-phospholipid interactions can modulate bilayer/non-bilayer transitions. Biochimica et Biophysica Acta (BBA)-Biomembranes, 601, 235–240. 10.1016/0005-2736(80)90528-3

DeLano, W. L. (2002). Pymol: An open-source molecular graphics tool. CCP4 Newsl. Protein Crystallogr, 40(1), 82–92.

Dos Santos Morais, R., Delalande, O., Pérez, J., Mouret, L., Bondon, A., Martel, A., Appavou, M.-S., Le Rumeur, E., Hubert, J.-F., & Combet, S. (2017). Contrast-Matched Isotropic Bicelles: A Versatile Tool to Specifically Probe the Solution Structure of Peripheral Membrane Proteins Using SANS. Langmuir, 33(26), 6572–6580. 10.1021/acs.langmuir.7b01369

Dürr, U. H., Soong, R., & Ramamoorthy, A. (2013). When detergent meets bilayer: birth and coming of age of lipid bicelles. Prog Nucl Magn Reson Spectrosc, 69, 1–22. 10.1016/j.pnmrs.2013.01.001

Eastman, P., Swails, J., Chodera, J. D., McGibbon, R. T., Zhao, Y., Beauchamp, K. A., Wang, L.-P., Simmonett, A. C., Harrigan, M. P., Stern, C. D., Wiewiora, R. P., Brooks, B. R., & Pande, V. S. (2017). OpenMM 7: Rapid development of high performance algorithms for molecular dynamics. PLOS Computational Biology, 13(7), e1005659. 10.1371/journal.pcbi.1005659

Eeman, M., & Deleu, M. (2010). From biological membranes to biomimetic model membranes. *Biotechnologie, Agronomie*, Société et Environnement, 14, 719–736.

Eichberg, J. (2002). Myelin P0: new knowledge and new roles. Neurochem Res, 27(11), 1331–1340. 10.1023/a:1021619631869

Fabrizi, G. M., Pellegrini, M., Angiari, C., Cavallaro, T., Morini, A., Taioli, F., Cabrini, I., Orrico, D., & Rizzuto, N. (2006). Gene dosage sensitivity of a novel mutation in the intracellular domain of P0 associated with Charcot-Marie-Tooth disease type 1B. Neuromuscul Disord, 16(3), 183–187. 10.1016/j.nmd.2006.01.006

Filbin, M. T., Zhang, K., Li, W., & Gao, Y. (1999). Characterization of the Effect on Adhesion of Different Mutations in Myelin P(0) Protein. Ann N Y Acad Sci, 883(1), 160–167. 10.1111/j.1749-6632.1999.tb08579.x

Gao, Y., Li, W., & Filbin, M. T. (2000). Acylation of myelin Po protein is required for adhesion. Journal of Neuroscience Research, 60(6), 704–713. 10.1002/1097-4547(20000615)60:6<704::AID-JNR2>3.0.CO;2-5

Giudice, R. D., Paracini, N., Laursen, T., Blanchet, C., Roosen-Runge, F., & Cárdenas, M. (2022). Expanding the Toolbox for Bicelle-Forming Surfactant–Lipid Mixtures. Molecules, 27(21), 7628. https://www.mdpi.com/1420-3049/27/21/7628 https://mdpi-res.com/d_attachment/molecules/molecules-27-07628/article_deploy/molecules-27-07628-v2.pdf?version=1667886204

Grandis, M., Vigo, T., Passalacqua, M., Jain, M., Scazzola, S., La Padula, V., Brucal, M., Benvenuto, F., Nobbio, L., Cadoni, A., Mancardi, G. L., Kamholz, J., Shy, M. E., & Schenone, A. (2008). Different cellular and molecular mechanisms for early and late-onset myelin protein zero mutations. Hum Mol Genet, 17(13), 1877–1889. 10.1093/hmg/ddn083

Grant, T. D. (2018). Ab initio electron density determination directly from solution scattering data. Nature Methods, 15(3), 191–193. 10.1038/nmeth.4581

Hägerbäumer, P., Gräbitz-Bräuer, F., Annegarn, M., Dargel, C., Stank, T. J., Bizien, T., & Hellweg, T. (2023). Interactions between DMPC Model Membranes, the Drug Naproxen, and the Saponin β-Aescin. Pharmaceutics, 15(2). 10.3390/pharmaceutics15020379

Halaby, D. M., & Mornon, J. P. (1998). The immunoglobulin superfamily: an insight on its tissular, species, and functional diversity. J Mol Evol, 46(4), 389–400. 10.1007/pl00006318

Hattori, N., Yamamoto, M., Yoshihara, T., Koike, H., Nakagawa, M., Yoshikawa, H., Ohnishi, A., Hayasaka, K., Onodera, O., Baba, M., Yasuda, H., Saito, T., Nakashima, K., Kira, J., Kaji, R., Oka, N., & Sobue, G. (2003). Demyelinating and axonal features of Charcot-Marie-Tooth disease with mutations of myelin-related proteins (PMP22, MPZ and Cx32): a clinicopathological study of 205 Japanese patients. *Brain*, *126*(Pt 1), 134-151. 10.1093/brain/awg012

Hong, H. (2014). Toward understanding driving forces in membrane protein folding. Archives of Biochemistry and Biophysics, 564, 297–313. 10.1016/j.abb.2014.07.031

Hopkins, J. B., Gillilan, R. E., & Skou, S. (2017). BioXTAS RAW: improvements to a free open-source program for small-angle X-ray scattering data reduction and analysis. J Appl Crystallogr, 50(Pt 5), 1545–1553. 10.1107/s1600576717011438

Hsu, N.-W., Nouri, B., Chen, L.-T., & Chen, H.-L. (2020). Hexagonal Close-Packed Sphere Phase of Conformationally Symmetric Block Copolymer. Macromolecules, 53(21), 9665–9675. 10.1021/acs.macromol.0c01445

Inouye, H., Ganser, A. L., & Kirschner, D. A. (1985). Shiverer and Normal Peripheral Myelin Compared: Basic Protein Localization, Membrane Interactions, and Lipid Composition. Journal of Neurochemistry, 45(6), 1911–1922. 10.1111/j.1471-4159.1985.tb10551.x

Inouye, H., & Kirschner, D. A. (2016). Evolution of myelin ultrastructure and the major structural myelin proteins. Brain Res, 1641(Pt A), 43–63. 10.1016/j.brainres.2015.10.037

Jean Harry, G., & Toews, A. D. (1998). Chapter 4 - Myelination, Dysmyelination, and Demyelination. In W. Slikker & L. W. Chang (Eds.), Handbook of Developmental Neurotoxicology (pp. 87–115). Academic Press. 10.1016/B978-012648860-9.50007-8

Jessen, K. R., & Mirsky, R. (1999). Schwann cells and their precursors emerge as major regulators of nerve development. Trends Neurosci, 22(9), 402–410. 10.1016/s0166-2236(98)01391-5

Jumper, J., Evans, R., Pritzel, A., Green, T., Figurnov, M., Ronneberger, O., Tunyasuvunakool, K., Bates, R., Žídek, A., Potapenko, A., Bridgland, A., Meyer, C., Kohl, S. A. A., Ballard, A. J., Cowie, A., Romera-Paredes, B., Nikolov, S., Jain, R., Adler, J., … Hassabis, D. (2021). Highly accurate protein structure prediction with AlphaFold. Nature, 596(7873), 583–589. 10.1038/s41586-021-03819-2

Kahnt, M., Klementiev, K., Haghighat, V., Weninger, C., Plivelic, T. S., Terry, A. E., & Björling, A. (2021). Measurement of the coherent beam properties at the CoSAXS beamline. Journal of Synchrotron Radiation, 28, 1948–1953. 10.1107/S1600577521009140

Katsaras, J., Harroun, T. A., Pencer, J., & Nieh, M.-P. (2005). “Bicellar” Lipid Mixtures as used in Biochemical and Biophysical Studies. Naturwissenschaften, 92(8), 355–366. 10.1007/s00114-005-0641-1

Kim, S., Jeon, T.-J., Oberai, A., Yang, D., Schmidt, J. J., & Bowie, J. U. (2005). Transmembrane glycine zippers: Physiological and pathological roles in membrane proteins. Proceedings of the National Academy of Sciences, 102(40), 14278–14283. doi:10.1073/pnas.0501234102

Kister, A., & Kister, I. (2023). Overview of myelin, major myelin lipids, and myelin-associated proteins [Review]. Frontiers in Chemistry, 10. 10.3389/fchem.2022.1041961

Krokengen, O. C., Raasakka, A., & Kursula, P. (2023). The intrinsically disordered protein glue of the myelin major dense line: Linking AlphaFold2 predictions to experimental data. Biochemistry and Biophysics Reports, 34, 101474. 10.1016/j.bbrep.2023.101474

Kulkarni, C. V., Wachter, W., Iglesias-Salto, G., Engelskirchen, S., & Ahualli, S. (2011). Monoolein: a magic lipid? [10.1039/C0CP01539C]. Physical Chemistry Chemical Physics, 13(8), 3004-3021. 10.1039/C0CP01539C

Laulumaa S, N. T., Raasakka A, Krokengen OC, Safaryan A, Hallin EI, et al. (2018). Structure and dynamics of a human myelin protein P2 portal region mutant indicate opening of the beta-barrel in fatty acid binding protein. BMC Structural Biology(18(1)).

Lemke, G., & Axel, R. (1985). Isolation and sequence of a cDNA encoding the major structural protein of peripheral myelin. Cell, 40(3), 501–508. 10.1016/0092-8674(85)90198-9

Li, M., Morales, H. H., Katsaras, J., Kučerka, N., Yang, Y., Macdonald, P. M., & Nieh, M.-P. (2013). Morphological Characterization of DMPC/CHAPSO Bicellar Mixtures: A Combined SANS and NMR Study. Langmuir, 29(51), 15943–15957. 10.1021/la402799b

Li, N., Perutkova, S., & Rappolt, M. (2017). My first electron density map: A beginner’s guide to small angle X-ray diffraction. Elektrotehniski Vestnik/Electrotechnical Review, 84.

Lin, S.-H., & Guidotti, G. (2009). Chapter 35 Purification of Membrane Proteins. In R. R. Burgess & M. P. Deutscher (Eds.), Methods in Enzymology (Vol. 463, pp. 619–629). Academic Press. 10.1016/S0076-6879(09)63035-4

Liu, Z., Wang, Y., Yedidi, R. S., Brunzelle, J. S., Kovari, I. A., Sohi, J., Kamholz, J., & Kovari, L. C. (2012). Crystal structure of the extracellular domain of human myelin protein zero. Proteins, 80(1), 307–313. 10.1002/prot.23164

Lombardo, D., Kiselev, M. A., Magazù, S., & Calandra, P. (2015). Amphiphiles Self-Assembly: Basic Concepts and Future Perspectives of Supramolecular Approaches. Advances in Condensed Matter Physics, 2015, 151683. 10.1155/2015/151683

Luo, X., Sharma, D., Inouye, H., Lee, D., Avila, R. L., Salmona, M., & Kirschner, D. A. (2007). Cytoplasmic domain of human myelin protein zero likely folded as beta-structure in compact myelin. Biophysical journal, 92(5), 1585–1597. 10.1529/biophysj.106.094722

Maeda, M. H., Mitsui, J., Soong, B.-W., Takahashi, Y., Ishiura, H., Hayashi, S., Shirota, Y., Ichikawa, Y., Matsumoto, H., Arai, M., Okamoto, T., Miyama, S., Shimizu, J., Inazawa, J., Goto, J., & Tsuji, S. (2012). Increased gene dosage of myelin protein zero causes Charcot-Marie-Tooth disease. Annals of Neurology, 71(1), 84–92. 10.1002/ana.22658

Manalastas-Cantos, K., Konarev, P. V., Hajizadeh, N. R., Kikhney, A. G., Petoukhov, M. V., Molodenskiy, D. S., Panjkovich, A., Mertens, H. D. T., Gruzinov, A., Borges, C., Jeffries, C. M., Svergun, D. I., & Franke, D. (2021). ATSAS 3.0: expanded functionality and new tools for small-angle scattering data analysis. J Appl Crystallogr, 54(Pt 1), 343–355. 10.1107/s1600576720013412

Mandich, P., Fossa, P., Capponi, S., Geroldi, A., Acquaviva, M., Gulli, R., Ciotti, P., Manganelli, F., Grandis, M., & Bellone, E. (2009). Clinical features and molecular modelling of novel MPZ mutations in demyelinating and axonal neuropathies. European Journal of Human Genetics, 17(9), 1129–1134. 10.1038/ejhg.2009.37

Marinko, J. T., Huang, H., Penn, W. D., Capra, J. A., Schlebach, J. P., & Sanders, C. R. (2019). Folding and Misfolding of Human Membrane Proteins in Health and Disease: From Single Molecules to Cellular Proteostasis. Chemical Reviews, 119(9), 5537–5606. 10.1021/acs.chemrev.8b00532

Martini, R., Zielasek, J., Toyka, K. V., Giese, K. P., & Schachner, M. (1995). Protein zero (P0)-deficient mice show myelin degeneration in peripheral nerves characteristic of inherited human neuropathies. Nat Genet, 11(3), 281–286. 10.1038/ng1195-281

Miltenberger-Miltenyi, G., Schwarzbraun, T., Loscher, W. N., Wanschitz, J., Windpassinger, C., Duba, H. C., Seidl, R., Albrecht, G., Weirich-Schwaiger, H., Zoller, H., Utermann, G., Auer-Grumbach, M., & Janecke, A. R. (2009). Identification and in silico analysis of 14 novel GJB1, MPZ and PMP22 gene mutations. Eur J Hum Genet, 17(9), 1154–1159. 10.1038/ejhg.2009.29

Min, Y., Kristiansen, K., Boggs, J. M., Husted, C., Zasadzinski, J. A., & Israelachvili, J. (2009). Interaction forces and adhesion of supported myelin lipid bilayers modulated by myelin basic protein. Proc Natl Acad Sci U S A, 106(9), 3154–3159. 10.1073/pnas.0813110106

Mirdita, M., Schütze, K., Moriwaki, Y., Heo, L., Ovchinnikov, S., & Steinegger, M. (2022). ColabFold: making protein folding accessible to all. Nature Methods, 19(6), 679–682. 10.1038/s41592-022-01488-1

Mittendorf, K. F., Marinko, J. T., Hampton, C. M., Ke, Z., Hadziselimovic, A., Schlebach, J. P., Law, C. L., Li, J., Wright, E. R., Sanders, C. R., & Ohi, M. D. (2017). Peripheral myelin protein 22 alters membrane architecture. Sci Adv, 3(7), e1700220. 10.1126/sciadv.1700220

Murugova, T. N., Ivankov, O. I., Ryzhykau, Y. L., Soloviov, D. V., Kovalev, K. V., Skachkova, D. V., Round, A., Baeken, C., Ishchenko, A. V., Volkov, O. A., Rogachev, A. V., Vlasov, A. V., Kuklin, A. I., & Gordeliy, V. I. (2022). Mechanisms of membrane protein crystallization in ‘bicelles’. Scientific Reports, 12(1), 11109. 10.1038/s41598-022-13945-0

Natali, F., Gliozzi, A., Rolandi, R., Relini, A., Cavatorta, P., Deriu, A., Fasano, A., & Riccio, P. (2002). Changes in the anisotropy of oriented membrane dynamics induced by myelin basic protein. Applied Physics A, 74(1), s1582–s1584. 10.1007/s003390201711

Nave, K.-A., & Boespflug-Tanguy, O. (1996). X-Linked Developmental Defects of Myelination: From Mouse Mutants to Human Genetic Diseases. The Neuroscientist, 2(1), 33–43. 10.1177/107385849600200111

Petoukhov, M. V., Franke, D., Shkumatov, A. V., Tria, G., Kikhney, A. G., Gajda, M., Gorba, C., Mertens, H. D., Konarev, P. V., & Svergun, D. I. (2012). New developments in the ATSAS program package for small-angle scattering data analysis. J Appl Crystallogr, 45(Pt 2), 342–350. 10.1107/s0021889812007662

Planté-Bordeneuve, V., Parman, Y., Guiochon-Mantel, A., Alj, Y., Deymeer, F., Serdaroglu, P., Eraksoy, M., & Said, G. (2001). The range of chronic demyelinating neuropathy of infancy: a clinico-pathological and genetic study of 15 unrelated cases. J Neurol, 248(9), 795–803. 10.1007/s004150170096

Plotkowski, M. L., Kim, S., Phillips, M. L., Partridge, A. W., Deber, C. M., & Bowie, J. U. (2007). Transmembrane Domain of Myelin Protein Zero Can Form Dimers: Possible Implications for Myelin Construction. Biochemistry, 46(43), 12164–12173. 10.1021/bi701066h

Poitelon, Y., Kopec, A. M., & Belin, S. (2020). Myelin Fat Facts: An Overview of Lipids and Fatty Acid Metabolism. Cells, 9(4). 10.3390/cells9040812

Prada, V., Passalacqua, M., Bono, M., Luzzi, P., Scazzola, S., Nobbio, L. A., Capponi, S., Bellone, E., Mandich, P., Mancardi, G., Shy, M., Schenone, A., & Grandis, M. (2012). Gain of glycosylation: a new pathomechanism of myelin protein zero mutations. Ann Neurol, 71(3), 427–431. 10.1002/ana.22695

Prosser, R. S., Evanics, F., Kitevski, J. L., & Al-Abdul-Wahid, M. S. (2006). Current Applications of Bicelles in NMR Studies of Membrane-Associated Amphiphiles and Proteins. Biochemistry, 45(28), 8453–8465. 10.1021/bi060615u

Purves, D., Augustine, G., Fitzpatrick, D., &, et al., e. (2001). Increased Conduction Velocity as a Result of Myelination. In Neuroscience (2nd ed.). Sinauer Associates. https://www.ncbi.nlm.nih.gov/books/NBK10921/

Quarles, R. H. (2002). Myelin sheaths: glycoproteins involved in their formation, maintenance and degeneration. Cell Mol Life Sci, 59(11), 1851–1871. 10.1007/pl00012510

Raasakka, A., Jones, N. C., Hoffmann, S. V., & Kursula, P. (2019a). Ionic strength and calcium regulate membrane interactions of myelin basic protein and the cytoplasmic domain of myelin protein zero. Biochemical and Biophysical Research Communications, 511(1), 7–12. 10.1016/j.bbrc.2019.02.025

Raasakka, A., & Kursula, P. (2020). How Does Protein Zero Assemble Compact Myelin? Cells, 9(8). 10.3390/cells9081832

Raasakka, A., Ruskamo, S., Barker, R., Krokengen, O. C., Vatne, G. H., Kristiansen, C. K., Hallin, E. I., Skoda, M. W. A., Bergmann, U., Wacklin-Knecht, H., Jones, N. C., Hoffmann, S. V., & Kursula, P. (2019b). Neuropathy-related mutations alter the membrane binding properties of the human myelin protein P0 cytoplasmic tail. PLOS ONE, 14(6), e0216833. 10.1371/journal.pone.0216833

Raasakka, A., Ruskamo, S., Kowal, J., Barker, R., Baumann, A., Martel, A., Tuusa, J., Myllykoski, M., Bürck, J., Ulrich, A. S., Stahlberg, H., & Kursula, P. (2017). Membrane Association Landscape of Myelin Basic Protein Portrays Formation of the Myelin Major Dense Line. Scientific Reports, 7(1), 4974. 10.1038/s41598-017-05364-3

Raasakka, A., Ruskamo, S., Kowal, J., Han, H., Baumann, A., Myllykoski, M., Fasano, A., Rossano, R., Riccio, P., Bürck, J., Ulrich, A. S., Stahlberg, H., & Kursula, P. (2019c). Molecular structure and function of myelin protein P0 in membrane stacking. Scientific Reports, 9(1), 642. 10.1038/s41598-018-37009-4

Rappolt, M., Hickel, A., Bringezu, F., & Lohner, K. (2003). Mechanism of the Lamellar/Inverse Hexagonal Phase Transition Examined by High Resolution X-Ray Diffraction. Biophysical journal, 84(5), 3111–3122. 10.1016/S0006-3495(03)70036-8

Robaglia-Schlupp, A., Pizant, J., Norreel, J. C., Passage, E., Sabéran-Djoneidi, D., Ansaldi, J. L., Vinay, L., Figarella-Branger, D., Lévy, N., Clarac, F., Cau, P., Pellissier, J. F., & Fontés, M. (2002). PMP22 overexpression causes dysmyelination in mice. Brain, 125(10), 2213–2221. 10.1093/brain/awf230

Ruskamo, S., Krokengen, O. C., Kowal, J., Nieminen, T., Lehtimäki, M., Raasakka, A., Dandey, V. P., Vattulainen, I., Stahlberg, H., & Kursula, P. (2020). Cryo-EM, X-ray diffraction, and atomistic simulations reveal determinants for the formation of a supramolecular myelin-like proteolipid lattice. Journal of Biological Chemistry, 295(26), 8692–8705. 10.1074/jbc.RA120.013087

Ruskamo, S., Raasakka, A., Pedersen, J. S., Martel, A., Skubnik, K., Darwish, T., Porcar, L., & Kursula, P. (2022). Human myelin proteolipid protein structure and lipid bilayer stacking. Cell Mol Life Sci, 79(8), 419. 10.1007/s00018-022-04428-6

Ruskamo, S., Yadav, R. P., Sharma, S., Lehtimäki, M., Laulumaa, S., Aggarwal, S., Simons, M., Bürck, J., Ulrich, A. S., Juffer, A. H., Kursula, I., & Kursula, P. (2014). Atomic resolution view into the structure-function relationships of the human myelin peripheral membrane protein P2. Acta Crystallogr D Biol Crystallogr, 70(Pt 1), 165–176. 10.1107/s1399004713027910

Schneider-Gold, C., Kötting, J., Epplen, J. T., Gold, R., & Gerding, W. M. (2010). Unusual Charcot-Marie-Tooth phenotype due to a mutation within the intracellular domain of myelin protein zero. Muscle Nerve, 41(4), 550–554. 10.1002/mus.21523

Sebinelli, H. G., Borin, I. A., Ciancaglini, P., & Bolean, M. (2019). Topographical and mechanical properties of liposome surfaces harboring Na, K-ATPase by means of atomic force microscopy [10.1039/C9SM00040B]. Soft Matter, 15(13), 2737–2745. 10.1039/C9SM00040B

Sedzik, J., Blaurock, A. E., & Hoechli, M. (1985). Reconstituted P2/myelin-lipid multilayers. J Neurochem, 45(3), 844–852. 10.1111/j.1471-4159.1985.tb04071.x

Shaharabani, R., Ram-On, M., Avinery, R., Aharoni, R., Arnon, R., Talmon, Y., & Beck, R. (2016). Structural Transition in Myelin Membrane as Initiator of Multiple Sclerosis. Journal of the American Chemical Society, 138(37), 12159–12165. 10.1021/jacs.6b04826

Shaharabani, R., Ram-On, M., Talmon, Y., & Beck, R. (2018). Pathological transitions in myelin membranes driven by environmental and multiple sclerosis conditions. Proc Natl Acad Sci U S A, 115(44), 11156–11161. 10.1073/pnas.1804275115

Shapiro, L., Doyle, J. P., Hensley, P., Colman, D. R., & Hendrickson, W. A. (1996). Crystal structure of the extracellular domain from P0, the major structural protein of peripheral nerve myelin. Neuron, 17(3), 435–449. 10.1016/s0896-6273(00)80176-2

Sharma, A., & Sharma, U. S. (1997). Liposomes in drug delivery: Progress and limitations. International Journal of Pharmaceutics, 154(2), 123–140. 10.1016/S0378-5173(97)00135-X

Shy, M. E. (2006). Peripheral neuropathies caused by mutations in the myelin protein zero. Journal of the Neurological Sciences, 242(1), 55–66. 10.1016/j.jns.2005.11.015

Shy, M. E., Jani, A., Krajewski, K., Grandis, M., Lewis, R. A., Li, J., Shy, R. R., Balsamo, J., Lilien, J., Garbern, J. Y., & Kamholz, J. (2004). Phenotypic clustering in MPZ mutations. Brain, 127(Pt 2), 371–384. 10.1093/brain/awh048

Sorour, E., MacMillan, J., & Upadhyaya, M. (1997). Novel mutation of the myelin P0 gene in a CMT1B family. Hum Mutat, 9(1), 74–77. 10.1002/(sici)1098-1004(1997)9:1<74::Aid-humu16>3.0.Co;2-m

Sreij, R., Dargel, C., Geisler, P., Hertle, Y., Radulescu, A., Pasini, S., Perez, J., Moleiro, L. H., & Hellweg, T. (2018). DMPC vesicle structure and dynamics in the presence of low amounts of the saponin aescin [10.1039/C7CP08027A]. Physical Chemistry Chemical Physics, 20(14), 9070–9083. 10.1039/C7CP08027A

Street, V. A., Meekins, G., Lipe, H. P., Seltzer, W. K., Carter, G. T., Kraft, G. H., & Bird, T. D. (2002). Charcot-Marie-Tooth neuropathy: clinical phenotypes of four novel mutations in the MPZ and Cx 32 genes. Neuromuscul Disord, 12(7-8), 643–650. 10.1016/s0960-8966(02)00021-4

Su, Y., Brooks, D. G., Li, L., Lepercq, J., Trofatter, J. A., Ravetch, J. V., & Lebo, R. V. (1993). Myelin protein zero gene mutated in Charcot-Marie-tooth type 1B patients. Proc Natl Acad Sci U S A, 90(22), 10856–10860. 10.1073/pnas.90.22.10856

Suresh, S., Wang, C., Nanekar, R., Kursula, P., & Edwardson, J. M. (2010). Myelin basic protein and myelin protein 2 act synergistically to cause stacking of lipid bilayers. Biochemistry, 49(16), 3456–3463. 10.1021/bi100128h

Svergun, D. I., Petoukhov, M. V., & Koch, M. H. (2001). Determination of domain structure of proteins from X-ray solution scattering. Biophysical journal, 80(6), 2946–2953. 10.1016/s0006-3495(01)76260-1

Tesei, G., & Lindorff-Larsen, K. (2023). Improved predictions of phase behaviour of intrinsically disordered proteins by tuning the interaction range [version 2; peer review: 2 approved]. Open Research Europe, 2(94). 10.12688/openreseurope.14967.2

Tesei, G., Trolle, A. I., Jonsson, N., Betz, J., Knudsen, F. E., Pesce, F., Johansson, K. E., & Lindorff-Larsen, K. (2024). Conformational ensembles of the human intrinsically disordered proteome. Nature, 626(8000), 897–904. 10.1038/s41586-023-07004-5

Tria, G., Mertens, H. D., Kachala, M., & Svergun, D. I. (2015). Advanced ensemble modelling of flexible macromolecules using X-ray solution scattering. IUCrJ, 2(Pt 2), 207–217. 10.1107/s205225251500202x

Ujwal, R., & Bowie, J. U. (2011). Crystallizing membrane proteins using lipidic bicelles. Methods, 55(4), 337–341. 10.1016/j.ymeth.2011.09.020

Uusitalo, M., Klenow, M. B., Laulumaa, S., Blakeley, M. P., Simonsen, A. C., Ruskamo, S., & Kursula, P. (2021). Human myelin protein P2: from crystallography to time-lapse membrane imaging and neuropathy-associated variants. The FEBS Journal, 288(23), 6716–6735. 10.1111/febs.16079

Wang, L., & Sigworth, F. J. (2010). Liposomes on a streptavidin crystal: a system to study membrane proteins by cryo-EM. Methods Enzymol, 481, 147–164. 10.1016/s0076-6879(10)81007-9

Wong, M. H., & Filbin, M. T. (1994). The cytoplasmic domain of the myelin P0 protein influences the adhesive interactions of its extracellular domain. J Cell Biol, 126(4), 1089–1097. 10.1083/jcb.126.4.1089

Wong, M. H., & Filbin, M. T. (1996). Dominant-negative effect on adhesion by myelin Po protein truncated in its cytoplasmic domain. The Journal of cell biology, 134(6), 1531–1541. 10.1083/jcb.134.6.1531

Wrabetz, L., Feltri, M. L., Quattrini, A., Imperiale, D., Previtali, S., D’Antonio, M., Martini, R., Yin, X., Trapp, B. D., Zhou, L., Chiu, S. Y., & Messing, A. (2000). P(0) glycoprotein overexpression causes congenital hypomyelination of peripheral nerves. J Cell Biol, 148(5), 1021–1034. 10.1083/jcb.148.5.1021

Yao, X., Fan, X., & Yan, N. (2020). Cryo-EM analysis of a membrane protein embedded in the liposome. Proceedings of the National Academy of Sciences, 117(31), 18497–18503. doi:10.1073/pnas.2009385117

