## Supplementary Information for "The cytoplasmic tail of myelin protein zero induces morphological changes in lipid membranes"

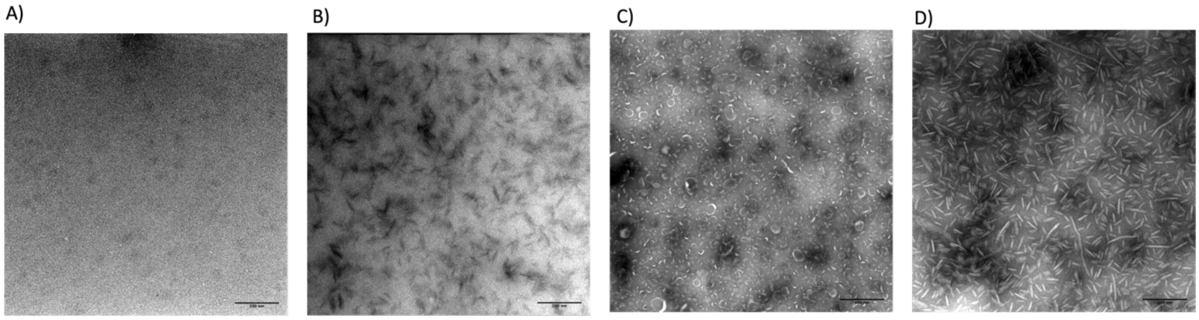

*Supplementary Figure 1: Control TEM images of A) HBS (20 mM HEPES, 150 mM NaCl pH 7.5), B) 20 uM P0ct, C) sonicated liposomes and D) bicelles with a q-ratio of 2.8.*

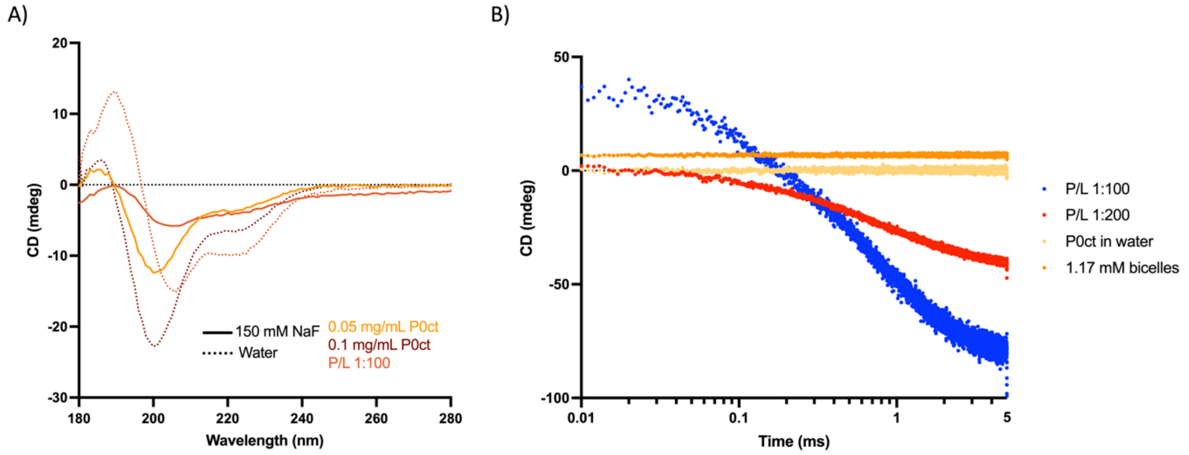

*Supplementary Figure 2: SRCD control experiments and time scale of P0ct-induced initial lipid turbidity. A) Control SRCD experiments for the stopped-flow kinetic measurements. Various concentration of P0ct and P/L 1:100 was measured with bicelles to verify secondary structures. Weak P/L 1:100 spectra with 150 mM NaF are likely due to heavy aggregations scattering light. B) Logarithmic time scale of rapid kinetics of P0ct-induced initial lipid turbidity. The SRCD signal was monitored at 195 nm for 5 s using stopped-flow measurements. P0ct was mixed with DMPC:DMPG (1:1) bicelles, q-ratio of 2.8 in two protein-to-lipid ratios. P0ct in solution and bicelles in solution were measured separately as controls.*

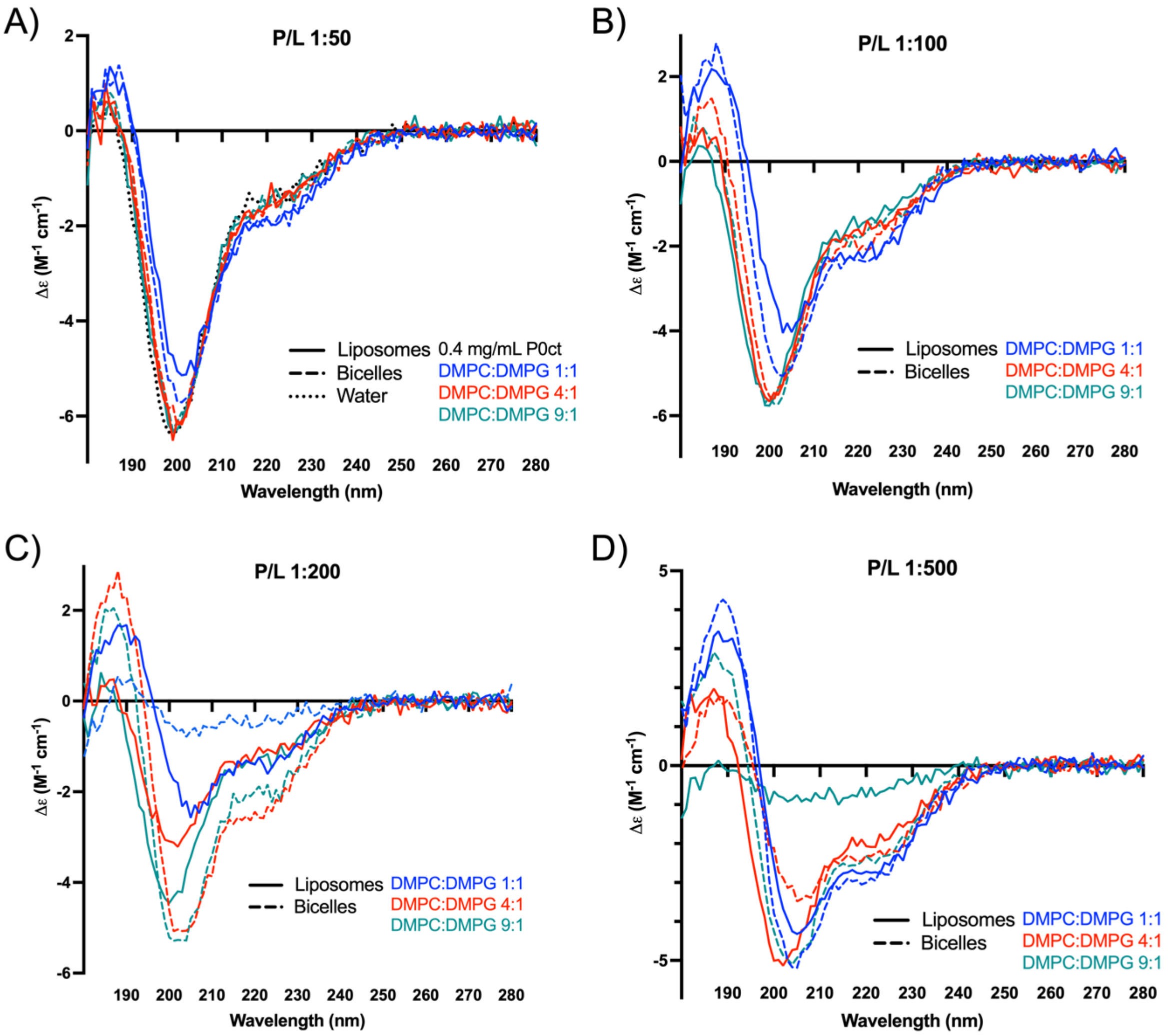

*Supplementary Figure 3: SRCD measurements of several protein-to-lipid ratios of P0ct in various DMPC:DMPG lipid mixtures with decreasing ratios of negatively charged lipids. The P0ct-liposomes/bicelle samples were all measured in water without any additives. Protein control in water is plotted in panel A for reference (dotted back line).*

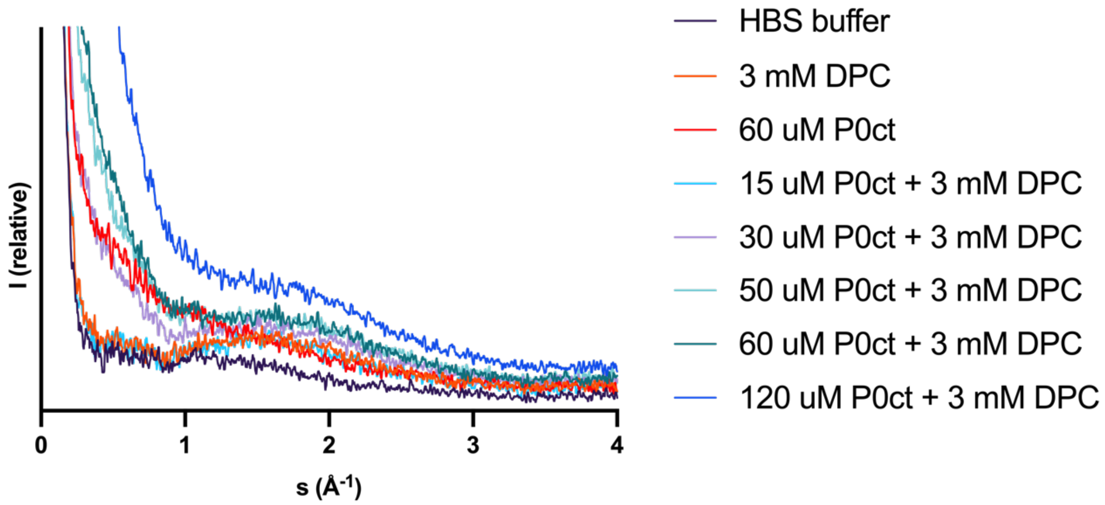

*Supplementary Figure 4: SAXD measurements of various protein-to-lipid ratios of P0ct and 3 mM DPC detergent. Different concentration of P0ct was added to 3 mM DPC to check for any potential diffraction peaks in P0ct-DPC systems. Buffer (20 mM HEPES, 150 mM NaCl, pH 7.5) and DPC were measured as controls.*

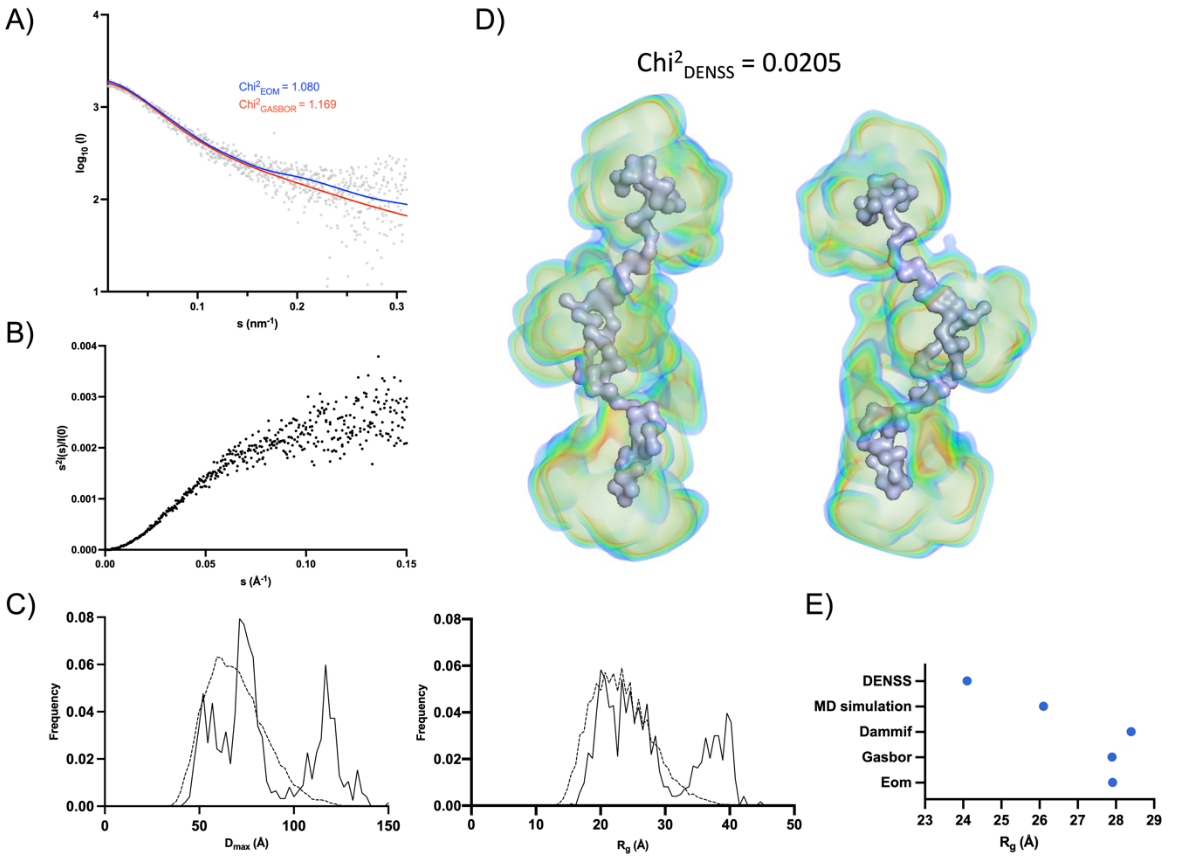

*Supplementary Figure 5: SEC-SAXS measurement and model of 14.4 mg/ml P0ct. A) Fit of top-ranked EOM and GASBOR model to the raw SAXS data. B) Kratky plot. C) $D_{max}$ and $R_g$ estimated by EOM. D) Highest-ranked GASBOR model of P0ct fitted inside the modelled electron density map by DENSS. The two models are related by a rotation of 180° for visualization. E) average $R_g$ from various modelling programs.*

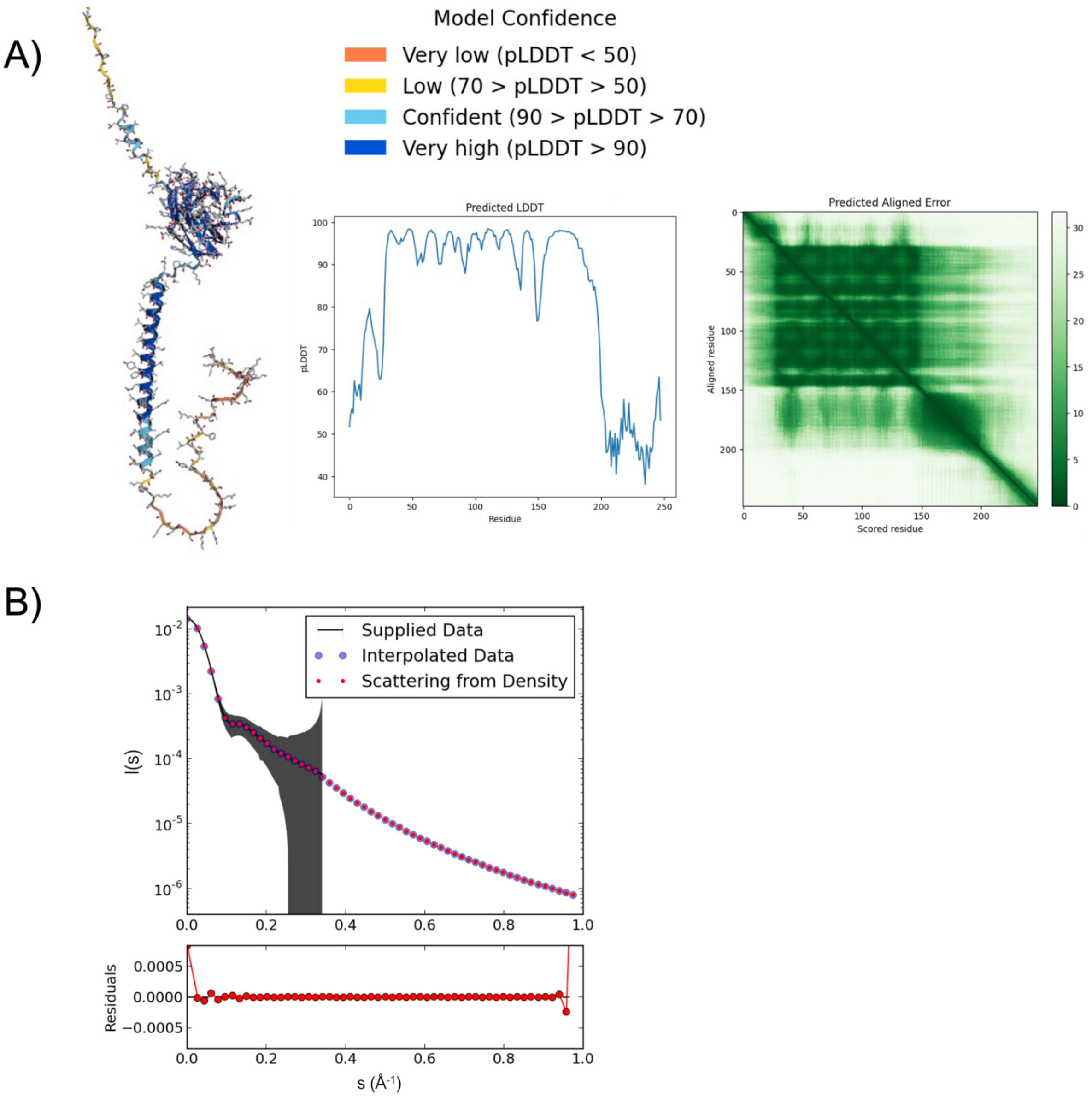

*Supplementary Figure 6: AlphaFold2 model and DENSS curve fit of full-length P0 (hP0). A) Highest ranked AlphaFold2 structure of P0ct colored by pLDDT, pLDDT plot and PAE plot. B) Fitting of the DENSS curve used for modelling the electron density map in DENSS.*

Supplementary Table 1: Non-linear fit parameters, two-phase decay, of the stopped-flow SRCD kinetics. Note that no change in signal was observed in the controls (P0ct and bicelles alone).

| | P/L 1:100 | P/L 1:200 | P0ct in water | Bicelles in water |
|---|---|---|---|---|
| **Two-phase decay** | | | | |
| **Best-fit values** | | | | |
| Y0 | 38.15 | 0.531 | 0.2233 | 6.852 |
| Plateau | -79.01 | -41.75 | -16.42 | -8.645 |
| PercentFast | 28.7 | 40.82 | Unstable | Unstable |
| KFast | 6.674 | 2.592 | ~ 0.0005250 | Unstable |
| KSlow | 0.9811 | 0.5886 | 0.000525 | 0.0003538 |
| Half Life (Slow) | 0.7065 | 1.178 | 1320 | 1959 |
| Half Life (Fast) | 0.1039 | 0.2674 | ~ 1320 | Unstable |
| Tau (slow) | 1.019 | 1.699 | 1905 | 2826 |
| Tau (fast) | 0.1498 | 0.3858 | ~ 1905 | Unstable |
| Rate constant ratio | 6.802 | 4.404 | ~ 1.000 | Unstable |
| **95% CI (profile likelihood)** | | | | |
| Y0 | 37.18 to 39.13 | 0.1727 to 0.8999 | 0.1894 to 0.2582 | 6.844 to 6.871 |
| Plateau | -79.19 to -78.83 | -42.17 to -41.42 | - | - |
| PercentFast | 27.67 to 29.79 | 36.74 to 45.37 | (Very wide) | (Very wide) |
| KFast | 6.113 to 7.277 | 2.320 to 2.908 | - | (Very wide) |
| KSlow | 0.9620 to 0.9991 | 0.5313 to 0.6385 | - | - |
| Half Life (Slow) | 0.6938 to 0.7205 | 1.086 to 1.305 | - | - |
| Half Life (Fast) | 0.09525 to 0.1134 | 0.2384 to 0.2988 | - | (Very wide) |
| Tau (slow) | 1.001 to 1.039 | 1.566 to 1.882 | - | - |
| Tau (fast) | 0.1374 to 0.1636 | 0.3439 to 0.4310 | - | (Very wide) |
| Goodness of Fit | | | | |
| Degrees of Freedom | 4995 | 9995 | 4995 | 4995 |
| R squared | 0.9892 | 0.9496 | 0.0004022 | 0.0005291 |
| Sum of Squares | 30055 | 44433 | 1974 | 590.7 |
| Sy.x | 2.453 | 2.108 | 0.6286 | 0.3439 |
| **Constraints** | | | | |
| PercentFast | 0 < PercentFast < 100 | 0 < PercentFast < 100 | 0 < PercentFast < 100 | 0 < PercentFast < 100 |
| KFast | KFast > 1*KSlow | KFast > 1*KSlow | KFast > 1*KSlow | KFast > 1*KSlow |
| KSlow | KSlow > 0 | KSlow > 0 | KSlow > 0 | KSlow > 0 |
| **Number of points** | | | | |
| # of X values | 10000 | 10000 | 10000 | 10000 |
| # Y values analyzed | 5000 | 10000 | 5000 | 5000 |

Supplementary table 2: SAXS parameters for the cytoplasmic domain of P0 (P0ct) and full-length P0 (hP0)

| **Data collection parameters** | **P0ct** | **hP0** |
|---|---|---|
| Instrument | CoSAXS, MAX IV laboratory | B21, Diamond Light Source |
| Wavelength (nm) | 0.1 | 0.1 |
| Angular range ($nm^{-1}$) | 0.001 - 0.3 | 0.0031 - 0.20 |
| Exposure time (s) | 0.200 | |
| Concentration (mg $ml^{-1}$) | 14.4 | 0.94 |
| Temperature (°C) | 25 | 20 |
| Frames collected | 4500 | 2567 |
| Injection volume (μL) | 40 | 45 |
| Flow rate (mL/min) | 0.700 | 0.075 |
| Column | S200 increase 10/300 | Superdex 200 3.2x300 |
| **Structural parameters** | | |
| $I_0$ (relative) [from Guinier] | 1928 | 0.015 |
| $R_g$ (nm) [From Guinier] | 2.62 | 3.90 |
| $R_g$ (nm) [From GASBOR] | 2.79 | 3.99 |
| $R_g$ (nm) [from EOM ensemble] | 2.35 | - |
| $R_g$ (nm) [from Dammif ensemble] | 2.84 | 3.93 |
| $R_g$ (nm) [from MD simulation] | 2.61 | 4.28 |
| $R_g$ (nm) [from DENNS modelling] | 2.41 | 4.23 |
| $D_{max}$ (nm) [from GASBOR] | 12.10 | 13.9 |
| $D_{max}$ (nm) [from EOM ensemble] | 7.96 | - |
| **Molecular mass analysis** | | |
| Molecular mass Mr (kDa) [Qp] | 8.24 | 94.4 |
| Molecular mass Mr (kDa) [MoW] | 8.66 | 113 |
| Molecular mass Mr (kDa) [Vc] | 9.01 | 124 |
| Molecular mass Mr (kDa) [Bayesian Inference] | 9.50 | 109 |
| Theoretical Mr from sequence (kDa) | 7.99 | 27.6 |
| **Software** | | |
| Primary data reduction | PRIMUS | PRIMUS |
| Data processing | PRIMUS | PRIMUS |
| *Ab initio* analysis | GASBOR | GASBOR |
| Conformational ensemble analysis | EOM | CORAL/CHARMM-GUI |
| Validation and averaging | PRIMUS | PRIMUS |
| Three-dimensional graphics representation | PyMOL | PyMOL |
| Electron density map | DENSS | DENSS |
| Protein modelling by sequence | AlphaFold2 | AlphaFold2 |

**Supplementary movies**

Patch experiments.

- Supplementary movie 1: patch experiment of 10 μM P0ct (as seen in Figure 3B)
- Supplementary movie 2: control experiments of patches
